# High resolution mapping reveals hotspots and sex-biased recombination in *Populus trichocarpa*

**DOI:** 10.1101/2022.05.10.491397

**Authors:** Chanaka Roshan Abeyratne, David Macaya-Sanz, Ran Zhou, Kerrie W. Barry, Christopher Daum, Kathy Haiby, Anna Lipzen, Brian Stanton, Yuko Yoshinaga, Matthew Zane, Gerald A. Tuskan, Stephen P. DiFazio

## Abstract

Fine-scale meiotic recombination is fundamental to the outcome of natural and artificial selection. Here, dense genetic mapping and haplotype reconstruction were used to estimate recombination for a full factorial *Populus trichocarpa* cross of seven males and seven females. Genomes of the resulting 49 full-sib families (N=829 offspring) were re-sequenced, and high-fidelity biallelic SNP/INDELs and pedigree information were used to ascertain allelic phase and impute progeny genotypes to recover gametic haplotypes. The fourteen parental genetic maps contained 1820 SNP/INDELs on average that covered 376.7 Mb of physical length across 19 chromosomes. Comparison of parental and progeny haplotypes allowed fine-scale demarcation of cross-over (CO) regions, where 38,846 CO events in 1,658 gametes were observed. CO events were positively associated with gene density and negatively associated with GC content and long terminal repeats. One of the most striking findings was higher rates of COs in males in 8 out of 19 chromosomes. Regions with elevated male CO rates had lower gene density and GC content than windows showing no sex bias. High-resolution analysis identified 67 candidate CO hotspots spread throughout the genome. DNA sequence motifs enriched in these regions showed striking similarity to those of maize, Arabidopsis and wheat. These findings, and recombination estimates, will be useful for ongoing efforts to accelerate domestication of this and other biomass feedstocks, as well as future studies investigating broader questions related to evolutionary history, perennial development, phenology, wood formation, vegetative propagation, and dioecy that cannot be studied using annual plant model systems.

## INTRODUCTION

Meiotic recombination shuffles genetic variation, and may bring together beneficial alleles or purge deleterious alleles to create more fit haplotypes (Felsenstein 1974). In the complete absence of recombination, deleterious mutations would accumulate faster than selection can remove them (Muller 1964) and such a scenario would leave fitness consequences at the mercy of infrequent reverse mutations. Therefore, meiotic recombination is selectively favored and increases the efficiency of adaptive evolution in finite populations in concert with a backdrop of mutation, genetic drift and selection (Hill and Robertson 1966; Felsenstein 1974). In a breeding context, recombination serves to dampen gains made by artificial selection and yet it benefits breeding programs by reducing linkage drag. Accordingly, genome-wide local recombination rates in both natural and structured populations will be under selective pressure stemming from the need to maintain a trade-off between reducing genetic load and reducing the immediate negative effects of recombination load due to disruption of favorable combinations of alleles (Charlesworth and Barton 1996).

Meiotic recombination is a highly regulated process that occurs within germline cells of sexually reproducing organisms. It is initiated with programmed DNA double stranded breaks (DSBs) during meiotic prophase I (Choi and Henderson 2015). The DSB repair process resolves these strand breaks as either cross-over (CO) or non-cross-over (NCO) events leading to recombined chromosomes or gene conversions, respectively (Wang and Copenhaver 2018). In the latter case, haplotype fidelity is maintained on either side of the breakpoint, whereas CO events lead to linkage between haplotypes derived from different chromatids.

Recombination rates may differ inter or intra-specifically (Smukowski and Noor 2011; Bauer *et al*. 2013), between sexes (Lenormand and Dutheil 2005), and even across genomic regions within the same individual (Slavov *et al*. 2012; Rodgers-Melnick *et al*. 2015; Gion *et al*. 2016; Kianian *et al*. 2018). In certain species such as mouse, yeast and *Caenorhabditis elegans*, there is a stable upper limit for CO counts per chromosome (CO homeostasis). However, such a trend has not yet been observed in plants where CO counts show a linear relationship with DSBs. As observed in maize and *Arabidopsis* a minimum of one CO per chromosome is obligatory and results in proper segregation of chromosomes during meiosis (Sidhu *et al*. 2015; Lambing *et al*. 2017). On a chromosomal scale, regions near centromeres and telomeres generally show a reduction of recombination, although such patterns are not clearly displayed in relatively shorter, acrocentric or telocentric chromosomes (Haenel *et al*. 2018). Correspondingly, COs are not independently and uniformly distributed across a genome, and display non-random spatial heterogeneity among extensively studied species where fine-scale recombination hotspots are interspersed throughout the genome (Kauppi *et al*. 2004). Hotspots are generally 1-10 kbps in size and have high probability of carrying a CO event, compared to background recombination or flanking cold regions suppressed for recombination. However, CO hotspot existence, size, and frequency within a genome may widely vary from species to species.

Hotspots are well established in humans and mice in which the most salient feature associated is enrichment of DNA binding sequences for the histone methyl transferase PRDM9 (Baudat *et al*. 2010). PRDM9 orthologs are absent in non-mammalian taxa, including arthropods, birds, fungi, and plants. Instead, proteins involved in DSB repair, including SPO11, REC8, RAD51, and MUS81 have been associated with recombination hotpots (He *et al*. 2017; Lambing *et al*. 2020). In various plant species recombination frequency is reportedly associated with localized genomic features related to chromatin compaction such as histone remodeling, DNA-methylation, gene content, and repeat content, thus implying accessibility of DNA for DSBs is a major deciding factor (Saintenac *et al*. 2011; Yelina *et al*. 2012; Shilo *et al*. 2015; Rodgers-Melnick *et al*. 2016; Darrier *et al*. 2017; He *et al*. 2017; Choi *et al*. 2018). CO interference is an additional factor that may control spacing between adjacent CO events and is responsible for punctate distribution of COs across the genome. This process is driven by the interference sensitive (Type-I) DSB resolution pathway and is considered the dominant form displayed by most plant species (Choi and Henderson 2015; Wang and Copenhaver 2018).

Recombination is suppressed within sex chromosomes and sex-determination regions (SDRs) of nascent sex chromosomes and provides a plausible mechanism by which sex-based characteristics are perpetuated (Lenormand 2003; Charlesworth *et al*. 2005). Apart from this, some species of animals and plants show sex-based differences in rates of recombination observed in autosomes (Burt *et al*. 1991; Lenormand and Dutheil 2005; Kong *et al*. 2010). This bias may range from complete absence of recombination in one sex (achiasmy) as in the case of Drosophila, or differential rates (heterochiasmy, Lenormand 2003) observed between males and females where either sex can have the higher rate (Bherer *et al*. 2017; Kianian *et al*. 2018). The differences in recombination could also be extended to spatial localization and broader patterns across the genome in some species (Zelkowski *et al*. 2019; Sardell and Kirkpatrick 2020). Although sex-based differences in recombination were established early on, definitive local features that explain the finer scale variation still need clarification. Differential rates of recombination at various hierarchical scales are under selection (Dapper and Payseur 2017), and identification of associated conserved sequence motifs, molecular markers and other localized genomic features could provide insights into mechanisms underlying adaptive trait variation (Penalba and Wolf 2020).

Much progress has been made toward revealing fine-scale genome-wide recombination patterns in plant species such as *Arabidopsis thaliana, Zea mays, Triticum aestivum*, and *Oryza sativa* largely due to the availability of large structured mapping populations or multi-generation breeding programs for these species (Rodgers-Melnick *et al*. 2015; Wang and Copenhaver 2018; Rowan *et al*. 2019; Lambing *et al*. 2020). However, these species represent a biased subset of plants confounded by high levels of inbreeding and/or long histories of domestication. Conversely, there is a dearth of such studies in undomesticated, primarily outbreeding perennial species such as forest trees, which represent the majority of biomass in terrestrial ecosystems (Neale and Kremer 2011; Silva - Junior and Grattapaglia 2015). Trees are subject to considerably different adaptive, developmental, and reproductive selective pressures (Petit and Hampe 2006) that could profoundly shape their recombination landscapes in comparison to classical plant model systems such as *Arabidopsis* or maize.

Black cottonwood (*Populus trichocarpa*) is an undomesticated, outbreeding, pioneer riparian species with moderate life span that is widely used as a model for basic and applied research on many aspects of tree biology. *Populus* is also a promising renewable feedstock for bioenergy and bioproducts given its rich genomic resources, short-rotation cycle and desirable lignocellulosic characteristics (Bradshaw *et al*. 2000; Tuskan *et al*. 2006; Sannigrahi *et al*. 2010; Porth and El-Kassaby 2015). As such, both commercial and ecological success converge on similar adaptive traits (e.g., adventitious rooting of stem and branches). At this juncture, selection favoring trait combinations both in managed or natural populations depend heavily on the recombination landscape. *Populus* is a genetically tractable model system with high-quality male and female reference genomes, and a range of other molecular resources that make it a robust model system to use for studying recombination patterns.

Fine-scale genome-wide patterns of recombination can be studied using population-based linkage disequilibrium (LD) estimates, but these are subject to vagaries of population demography, drift, natural selection, and mutation rate, and they only provide information on sex averaged recombination rates (Auton and McVean 2007). Conversely, resolution of pedigree-based genetic maps is largely limited only by the size of the mapping population and the power of the genetic marker system to detect recombination breakpoints. Pedigrees can therefore provide insight into sex-based differences in recombination rates as well as more accurate estimates of single generation recombination rates. Most existing pedigree based genetic maps for *Populus* have been constructed using interspecific hybrids and large full-sib families and a limited number of molecular markers (Bradshaw *et al*. 1994; Yin *et al*. 2004; Gaudet *et al*. 2008). Recent advances in high-throughput whole-genome re-sequencing platforms, improved variant discovery methods and better reference genome assemblies facilitate the use of genome-wide SNP and INDEL markers at sufficient density to enable fine-scale mapping (Fang *et al*. 2018). However, mapping population size can still be a challenge, limiting the number of observed recombination events and map resolution.

Here we use a large-scale full-factorial cross in *P. trichocarpa*, coupled with whole-genome resequencing to reveal the genome-wide recombination landscape and patterns of inheritance at a finer scale than has previously been possible in undomesticated plants. We produced dense genetic maps for fourteen parents that contain a balanced representation of each sex. Data contributed from multiple individuals allowed us to conduct fine-scale analyses of sex dimorphism and intraspecific variation of genome-wide recombination patterns. We show how recombination rates vary within and among genomes, and between the sexes, elucidate key genomic features that may play a role in shaping recombination rates at a scale of approximately 960 kb.

## MATERIALS & METHODS

### *Populus trichocarpa* mapping population

The *P. trichocarpa* mapping population used in the study consists of a total of 829 progeny from 49 full-sib families derived from a full-factorial cross between seven males and seven females (Harman-Ware *et al*. 2021) (Table S1). The parents were originally collected as part of a population of 448 genotypes from natural riparian stands in WA and OR, USA (Figure S1; Wegrzyn *et al*. 2010). The parents were selected to represent the full range of natural variation in lignin composition observed in the population.

### DNA isolation, whole-genome resequencing and variant calling pipeline

Genomic DNA was extracted from foliage from all progeny and parents using the DNeasy 96 Plant DNA isolation kit (Cat. No. 69181; Qiagen, Valencia, CA, USA). The sample library preparation, quality control and whole-genome resequencing up to an expected coverage of 5× and 15× for all progeny and parental clones respectively, was carried out using the Illumina HiSeq 2500 and NovaSeq 6000 sequencing platforms subject to established standard quality control and sequencing protocols (https://jgi.doe.gov/user-programs/pmo-overview/protocols-sample-preparation-information/) at the DOE Joint Genome Institute (JGI). Paired-end reads were aligned to a male *P. trichocarpa* Stettler-14 male reference assembly (Hofmeister *et al*. 2020), modified as described in Zhou *et al*. (2020). Mapping was accomplished with BWA v.0.7.10-r789 with default parameters. The MarkDuplicates tool in Picard v.1.131 was used to locate and flag duplicate reads in BAM files. We used an adaptation of the short variant discovery best practices pipeline for Genome Analysis Toolkit v.4.2.0 (Depristo *et al*. 2011) to identify genome-wide SNP and INDELs. GATK’s HaplotypeCaller tool was used to call SNPs and INDELs simultaneously. A “truth dataset” containing 1,030,941 biallelic SNPs and 125,772 INDELs was derived from a starting set of 37,661,220 variants, by imposing strict biological constraints on Mendelian segregation and Mendelian violations (Table S2). The truth set was subsequently used to train GATK’s VQSR pipeline in order to identify 14,656,297 bi-allelic SNPs and INDELs enriched for true variants.

### Parental genotypes: error correction and phasing

Parentage was verified and corrected as needed for all offspring based on degree of match to putative parents using 1,731,769 hard-filtered SNPs that represented a random subset of genome-wide variants (Table S2). Parental genotypes were then imputed or corrected based on maximizing the likelihood of observing the validated half-sib family pedigree genotypes (Wang 2004). Given the moderate rate of missing genotypes in the offspring, estimations of joint-likelihood for predicted parental genotypes were based on at least a half-sib family size of 50 progeny.

A consensus of two independent methods was used to phase the parental chromosomes. Our first method assumed congruence in physical marker order between the focal parent and the reference genome, where pairwise linkage disequilibrium (LD) estimates between neighboring markers were used to incrementally infer parental haplotype configuration (i.e., positive LD meaning markers are in coupling phase and negative meaning repulsion phase). A semi-automated inspection was carried out to solve situations where the absolute value of LD between adjacent markers was low, meaning that the reference genome was inaccurate. As a precursor to our second independent method, corrected trio genotypes were used to identify the focal parent allele contribution in each offspring. Here if all individuals in the trio were heterozygous or Mendelian violations were observed, the focal parent allele contribution was designated as missing and loci with more than 25% missing data were excluded. The two-point pairwise linkage analysis implemented in the Onemap R-package v2.1.1 (Margarido *et al*. 2007) was used to cluster markers into phased linkage groups using the following cutoffs: max.rf = 0.5 and LOD > 8. The resulting framework haplotypes were then used to infer the phases of all intervening markers segregating in the family. The final parental haplotypes resulted from the consensus of both adjacent marker LD and Onemap based clustering approaches and consisted of approximately 900,000 fully-informative biallelic SNPs and INDEL markers for each parent.

### Offspring genotypes: phasing, imputation and demarcation of cross-over (CO) break-points

Initial offspring phasing was performed by identifying haplotypes derived from the common parent of each of the fourteen half-sib families (Figure S2). These haplotypes were refined by imputing offspring haplotype configurations in bins using all available segregating markers, based on a sliding-window smoothing algorithm that assumes CO events are rare within an LG. Each window contained 20 heterozygous markers, with 10 markers overlapping with neighboring windows. This smoothing algorithm was implemented from both ends of each LG to obtain the consensus. However, inferred CO region (i.e., the region between two markers where a CO could be unequivocally ascertained) resolution was limited by the number of high-quality informative markers in the flanking regions. The median size of the inferred CO regions was 30.71 kb (Figure S3). The minimum haplotype blocks were defined as regions exceeding 500 kb in size (excluding the ends of chromosomes). With this filtering, we are confident that we detected the vast majority of the COs for this mapping population while eliminating most gene conversion events.

### Construction of fourteen genetic linkage maps

A linkage map was constructed for the common parent of each half-sib family using the OneMap R-package. The marker datasets consisted of approximately, 900,000 fully-informative biallelic SNPs and INDEL markers spread across the whole genome for a given half-sib family. Since the parental haplotype configurations were fully determined, we recoded the markers for the common parent as an F1-backcross. Markers with completely redundant genotype information across offspring were binned for computational efficiency. Using the criteria of max.rf. = 0.5 and LOD > 8 yielded exactly 19 LGs per parent, corresponding to the base chromosome number of *Populus*. Marker order within LGs was determined by minimizing the total number of recombination events, implemented with the RECORD algorithm using Onemap’s “order_seq” function (Van Os *et al*. 2005) with the following adjustments of default parameters: n.init=8, subset.search=“twopt”, twopt.alg=“rec”, THRES=3, touchdown=TRUE. Markers that did not initially enter the framework map were tried again using ‘try_seq’ function with a THRES=2 to identify their most likely positions. Genetic distances between markers were estimated using the “Kosambi” mapping function allowing for possible CO interference. However, since a given marker order produced for each linkage group was not based on an exhaustive search, markers were rippled with a local window size of 4 using the “ripple_seq” function to identify and reposition misplaced markers using default parameters (ws=4, LO =3, tol=10e-5).

### Statistical models for analyzing chromosome scale sex-based differences in CO counts

To test the role of sex in determining CO counts on a chromosome scale, we used a generalized linear mixed model (GLMM) from the lme4 R-package and a Poisson link function as detailed below.

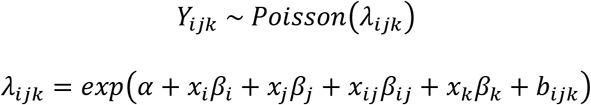

where, Y represents the total CO count and *i* =gender of the focal parent in the half-sib family (either female or male); *j* = chromosome identity (Chr01, Chr02,…, Chr19); *k* = identity of the half-sib family nested within i^th^ sex; λ = expected total CO count. β_i_, β_j_, β_ij_ and β_k_ are fixed effect coefficients for sex (female or male), chromosome identity, sex-chromosome interaction effect and halfsib family size, respectively, while x_i_, x_j_, x_ij_ and x_k_ denote their respective indicator variables. The last term, b_i|k_ represents half-sib family identity as a random factor nested within sex. The selected model provided the lowest Akaike Information Criterion (AIC) out of all models considered (Table S3). There were no discernible patterns in Pearson-residuals across any of the covariates used in this model or fitted values (Figure S4).

### Comparing average pairwise marker LD to cumulative CO counts

In order to analyze historic recombination rates in a natural population of *P. trichocarpa*, we used a collection of 220 unrelated trees from the portion of the *P. trichocarpa* range that overlaps with the collection sites of the parents of the 7×7 trial (Figure S1). The rationale for restricting the collection to this area was to avoid genetic structure that could have distorted linkage disequilibrium patterns. This subset of the population was identified from a larger collection that had been fully resequenced, as described in Chhetri *et al*. (2019). The resequenced reads for this natural population were aligned to the same Stettler-14 male reference genome that was used for the mapping population. An input variant dataset (SNPs) was derived using similar variant calling pipeline and parameters as described previously. The subpopulation was selected by evaluation of cluster admixture proportions using the software fastStructure (Raj *et al*. 2014) with the SNP dataset as inputs. Briefly, ten iterations were run with cluster number (K) from 1 to 7. The number of clusters was selected using the algorithm recommended by fastStructure developers and implemented in the script chooseK.py. Five clusters were the optimum to explain structure, one of them corresponding to the core population.

Assignment to that cluster was determined when the mean admixture proportion (Q value) of an individual for that cluster was above 0.8. In order to estimate LD within non-overlapping genomic windows, a subset of 13,989,405 SNPs was extracted for this subpopulation that intersected with the 7×7 SNP dataset, and this was used to estimate the squared correlation coefficient between (*r*^2^) genotypes using the vcftools -geno-r2 function with ld-window-bp parameter set to 10,000. The average pairwise LD was then calculated for the same set of non-overlapping genomic windows of 960 kbps that were used for the CO counts.

### Statistical model for analyzing finer-scale sex-based differences in cumulative CO counts

The finer scale analysis for heterochiasmy seeks to identify genomic regions that show differences in CO counts between sexes. As informed by the wavelet analysis (Supplementary methods), most of the variance in CO count signal for a given sex was contributed by lower dyadic scales (Figure S5 & S6). Yet, the dataset may suffer from poor resolution at finer scales such as 60 kb through 240 kb, largely due to the resolution of the CO region demarcation. At higher scales such as 3.84 Mb to 7.68 Mb, the analysis may be redundant with the chromosome scale analysis described above. The wavelet analysis identified a non-overlapping window size of 960 kb to be the most appropriate finer scale that unravels sex differences in CO rates. The response variable included cumulative CO counts for each sex at each non-overlapping window across the whole genome. First, windows that overlap putative centromeres and telomeres were removed by imposing a chromosome-wide minimum average CO count after determining the global distribution of CO counts across the genome. Windows from the tails of the index of dispersion distribution (≤ 5^th^ and ≥ 95^th^ percentile) were removed as outliers, resulting in a total of 280 windows to be tested. The effect of sex for each window was tested using a Poisson exact test implemented in the exactci R-package (Fay 2010), with mid-p-values (Heller and Gur 2011), and significance was determined using a false discovery procedure (FDR_BH_) as implemented in the p.adjust function in R (Benjamini and Hochberg 1995).

### Estimation of repeat content, gene content and AT/GC composition within genomic windows

Repeats were identified using the RepeatModeler (v1.0.8) package, and used to mask the genome with Repeatmasker v4.0.3 (Smit *et al*. 2013-2015) following the same approach as Zhou *et al*. (2020). The AT/GC composition, long terminal repeat (LTR) counts (i.e., *Gypsy, Copia* and simple repeats) and gene content were estimated in non-overlapping 960kb windows with bedtools utilities v2.17.0 (Quinlan and Hall 2010). Given that LTRs are harbored near telomeric and centromeric regions and observation that cumulative CO counts tend to show extreme values near these regions, genomic windows overlapping putative centromeric and telomeric regions were removed as outliers using a genome-wide fixed cutoff (cumulative CO count per window < 50).

### Analyzing association of genomic correlates and male biased heterochiasmy

AT/GC composition, LTR repeat counts, and gene content were estimated within non-overlapping 960-kb genomic windows as explained above. A subset (140) of the total genomic windows of the same size (960 kb), were identified in two comparator groups as male biased (15) and background (125). ‘Male biased windows’ included the 15 genomic windows that showed significant elevation of CO counts in males in comparison to females (described earlier in methods). A ‘background window’ was defined as one which exhibits a nominal male bias (female:male ratio < 1 at 0.4 ≤ FDR_BH_ ≥ 0.8). The association between each of the genomic correlates and type of window categorized as either ‘male biased’ or ‘background’ were investigated using a two tailed Wilcoxon rank-sum test.

### Localized CO pattern prediction using genomic correlates and sex

Genomic correlates were fitted to CO counts within genomic windows of size 960 kb, using backward stepwise linear regression with the ‘lm’ function in R (R Core Team 2013). All covariates were centered and scaled for this analysis and the following models were selected based on lowest AIC and homoscedasticity of standardized residuals (Table S4):

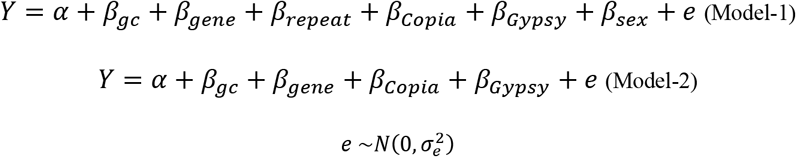

where, *Y* represents the total CO count for a half-sib family within a window; α, β_gc_, β_gene_, β_repeat_, β*_Copia_*, β*_Gypsy_* and β_sex_ are the intercept and fixed effects for percent GC content, count of genes, count of simple repeats, count of *Copia-LTR* elements, count of *Gypsy*-LTR elements, and binary factor of sex (either male of female) respectively, within a 960-kb non-overlapping genomic window; *e* represents the residual which is a random variable that is normally distributed with a mean of zero and variance σ^2^_e_. Model robustness of both models were assessed using 5-fold repeated cross-validation using the caret v 6.0-86 R-package (Kuhn 2008).

### Genome-wide CO hotspot identification

The genome was partitioned into 30-kb non-overlapping windows and each CO event was assigned to a single window based on the CO region mid-point. Window size was determined based on the median CO region (approximately 37 kb). CO hotspots were defined as windows showing significant deviation from the expected number of cumulative COs per window, with expectation under the null hypothesis that the probability of COs across the genome follows a Poisson distribution (λ=2.98; FWER_Bonferroni_ ≤ 0.05). Although a naïve approximation, stringent FWER cut-off enables detection of contiguous windows that far exceed the genome-wide expectation, highlighting candidate genomic regions enriched for CO hotspots. Due to structural rearrangements observed in parents GW-2393 and GW-6909, a stretch of windows was excluded from further analyses (more details in Results). Windows with < 2 informative markers on average within 60 kbps of window borders were removed to minimize the possibility of undetected COs in a neighboring window elevating the counts within a window. Also, windows where one or more parents presented anomalously high CO counts were removed as outliers. These outlier windows were flagged when index of dispersion for a given window (calculated using cumulative CO counts in half-sib families) exceeded the top 90^th^ percentile. Using similar methods, CO hotspots were also estimated separately for males and females (λ_female_ = 1.38; λ_male_ = 1.54; FWER_Bonferroni_ ≤ 0.05).

### Identification of DNA sequence motifs and genomic correlates associated with COs

Of the observed CO regions, 2054 that were demarcated to a 10-kb genomic region or less were used for this analysis. DNA sequence information in FASTA format for these narrowly demarcated CO regions were extracted using an in-house developed Perl script. These sequences were provided as a single set of positive inputs to the STREME (Simple, Thorough, Rapid, Enriched Motif Elicitation) algorithm within MEME v5.3.0 suite of software (Bailey *et al*. 2009), where differential enrichment to an automatically generated control set was investigated by setting the ‘--objfun’ parameter to ‘de’. The enriched DNA sequence motifs identified in this analysis were compared against a database of genomewide *15*-mers, by aligning DNA sequences in the two data sets to assess similarity. The genomewide *15*-mers and their counts were estimated for the Stettler-14 reference genome using jellyfish 2.2.10 (Marcais and Kingsford 2011). Enriched DNA sequence motifs were aligned to the *15*-mer database using blastn algorithm with modifications to standard settings (-dust no; -task blastn-short) on BLAST 2.3.0 (Altschul *et al*. 1990). In order to assess the similarity of CO associated DNA sequence motifs identified in this study, with such motifs identified in selected set of previous studies (Table S11), motifs were aligned to a second database consisting of DNA sequence motifs associated with recombination hotspots in *Arabidopsis*, maize and wheat using the same methods as explained earlier.

In order to investigate whether there are DNA sequence motifs enriched for CO hotspots in comparison to regions of the genome that show average CO counts, a second set of 119 non-overlapping 30-kb genomic regions previously identified as CO hotspots were provided as inputs to MEME. Similar parameters were used as before with the exception of now explicitly defining a negative control in the form of 3,214, 30-kb genomic regions that displayed average CO rates.

In order to investigate whether there are genomic features enriched for CO hotspots in comparison to genome-wide background levels, the same set of background genomic windows (3,124) were contrasted with 67 non-overlapping 30-kb genomic regions previously identified as CO hotspots. The full set of genomic correlates, AT/GC composition, LTR repeat counts (*Gypsy*, *Copia* and simple repeats) and gene content were all re-estimated for these 30-kb genomic windows and differential enrichment investigated using a two tailed Wilcoxon rank sum test.

## RESULTS

### Construction of 14 genetic linkage maps using marker segregation within half-sib families

All fourteen genetic-linkage maps (seven for each sex) converged on 19 linkage groups, representing the haploid chromosome count for *P. trichocarpa* (Table S5). This study represents the first attempt at producing multiple linkage maps for each sex using half-sib family marker segregation in a single *P. trichocarpa* mapping population. Framework genetic maps contained 1820 SNP/INDEL markers on average and consist exclusively of high-quality markers mapping to a single position with a conservative threshold. Genetic maps span 376.7 Mb of physical length across 19 chromosomes in the Stettler-14 male reference assembly (Table S6). Overall median physical distance between adjacent mapped markers was approximately 126 kbps and the median genetic distance was 1.03 cM. The genetic maps show high collinearity with the Stettler-14 male reference sequence assembly (**Figure 1)**. One notable exception is LG-XVII in parents GW-2393 & GW-6909, which show anomalously high map distances at the fringe of genomic region 9,044,429 bps to 10,304,428. This is likely an artifact caused by large structural rearrangements in these two parent genomes. Also, LG-I of GW-6909 was trimmed due to a lack of informative markers beyond 29 Mb due to a preponderance of very high segregation distortion in this region. These two regions were excluded from further chromosome-scale analyses to avoid possible biases due to structural artefacts.

**Figure 1:**
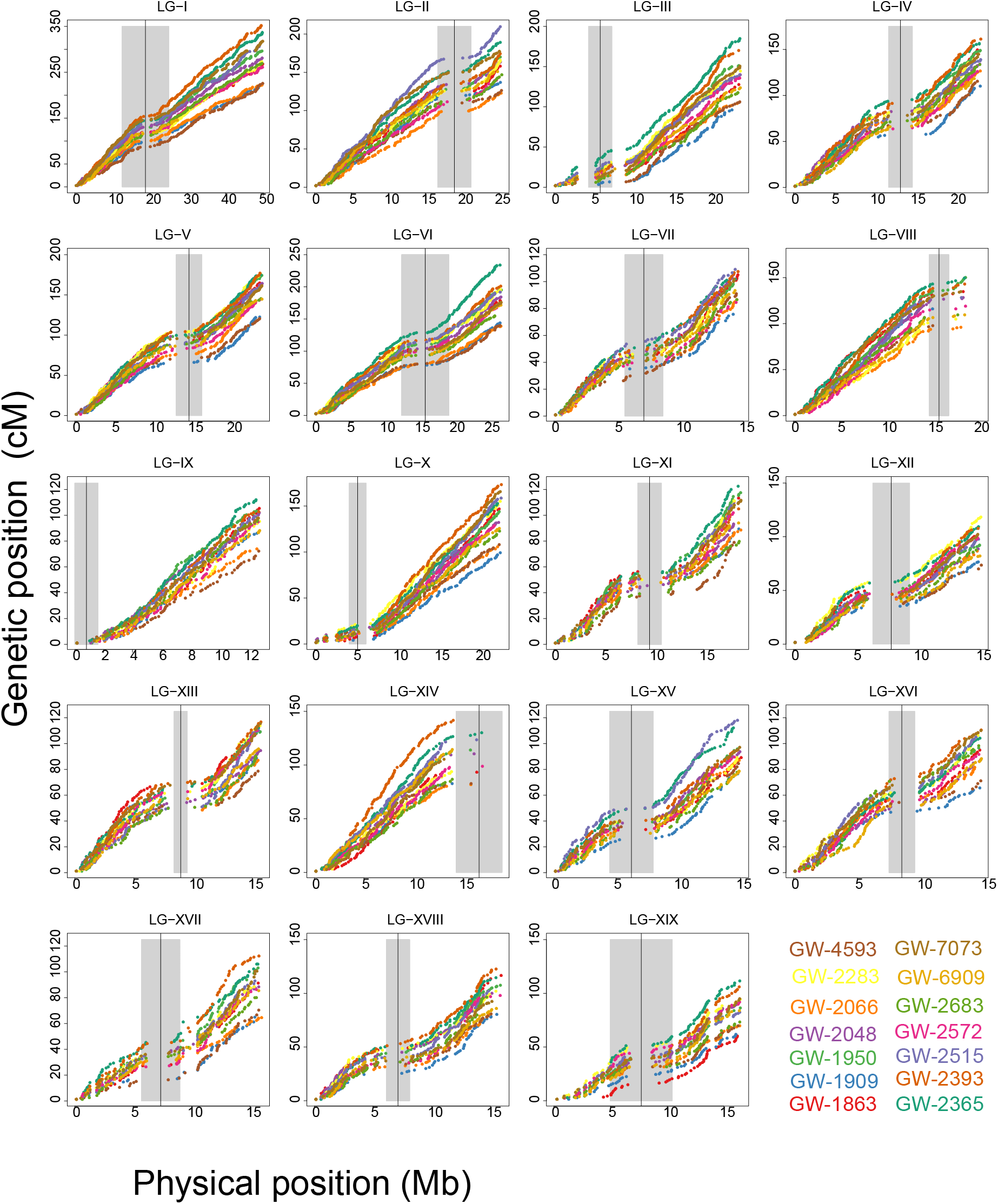
Fourteen linkage maps constructed using phased markers for each half-sib family. Genetic maps are presented as scatter plots of genetic versus physical map positions for all chromosomes. Each half-sib family is represented in a different color for each LG. Grey shading indicates the putative centromere boundaries. The Y-axis has been scaled to the same height across all scatter plots in this image.

Although parental maps generally show high collinearity of marker order, there is considerable variation in recombination rate among individuals, as demonstrated by divergence of the scatter-plots representing genetic maps of each half-sib family (**Figure 1**). Nevertheless, a high correlation (Figure S7) was observed between genetic map averages produced in our study in comparison to similar maps from an independent study using SSR markers (Yin *et al*. 2004). This was true for both male and female median map sizes (median female maps: *r* = 0.92, p = 1.709 x 10^-8^; median male maps: *r* = 0.93, p = 5.868 x 10^-9^). Average recombination rate across LGs fluctuated within a narrow range and varied from 4.24 – 7.53 cM·Mb^-1^ for females and 5.61 – 7.75 cM·Mb^-1^ males. The lowest and highest recombination rate estimates for each sex occurred on LG-XIX and LG-IX respectively (Table S6).

### Genome-wide recombination rate analyzed using cross-over (CO) counts

The accuracy of map-based estimates of recombination rate depends on informative marker density for the region under investigation. The median marker distance in our genetic maps was 126 kb. To improve resolution, we directly estimated CO rate by demarcating recombination breakpoints to a median resolution of 37 kb based on inferred haplotypes in the progeny. A total of 38,846 CO events were observed within 1658 gametes (829 diploid offspring) averaging approximately 1.2 COs per chromosome. Of the total observed, 18,355 and 20,491 COs occurred within female and male groups, respectively, indicating a significant gross cumulative difference of 2,136 CO events between the sexes (χ^2^_df=1_; p= 2.2×10^-16^).

### Chromosome-scale differences in CO rate between sexes

As expected LG identity was the most significant factor affecting CO count, and physical length of the LG explained much of the variance between the average number of COs within half-sib families (*r*^2^ = 0.77, p = 2.2×10^-16^) (Figure S8). Nevertheless, there was still a considerable amount of unexplained residual variance among half-sib families for a given LG. We observed unequal cumulative CO counts among parents of the two sexes within LGs I, II, III, VI, X, XIV, XVI and XIX (χ^2^_df=1_; FWER ≤ 0.05) which hinted at possible sex-based recombination rate differences (Table S7). Male parents showed higher CO rates (male-biased heterochiasmy) for 8 out of 19 LGs, but no LGs showed female-biased heterochiasmy. A generalized linear mixed model (GLMM) was used to model cumulative CO count data across half-sib families, that included sex as a fixed effect among other covariates (Table S8). This model yielded a significant positive coefficient for males, that translates to approximately 20% more COs in males in comparison to females. Furthermore, we observed that sex-based differences depended on the LG considered. The significant negative estimates for ‘LG × SEX’ interaction terms for LGs IV, V, VII, VIII, IX, XI, XII, XIII, XV, XVIII indicated reduced or absent male-biased heterochiasmy, which was consistent with lack of significant differences for these groups in the χ^2^ analysis of sex effects (Table S7).

### Wavelet analysis for identifying the scale of sex differences in CO counts

A fine-scale comparison of CO counts between males and females requires choosing a window size that minimizes spatial autocorrelation, while still providing sufficient resolution to detect underlying factors that drive differences between them. We implemented a wavelet analysis, which is a signal processing technique that can decompose the total variance in CO counts across a chromosome for a wide range of scales. This revealed the optimum scale at which features such as CO counts and associated genomic features could be compared. CO count differences between males and females were most prominent at scales 480 kb to 1.9 Mb. At finer scales, differences remained indiscernible and higher scales are redundant with the chromosome scale analysis presented earlier (**Figure 2; Figure S6**). The optimum window size was selected as 960 kb (Supplementary methods). All subsequent analysis related to feature contrasts between sexes are carried out at this scale.

**Figure 2:**
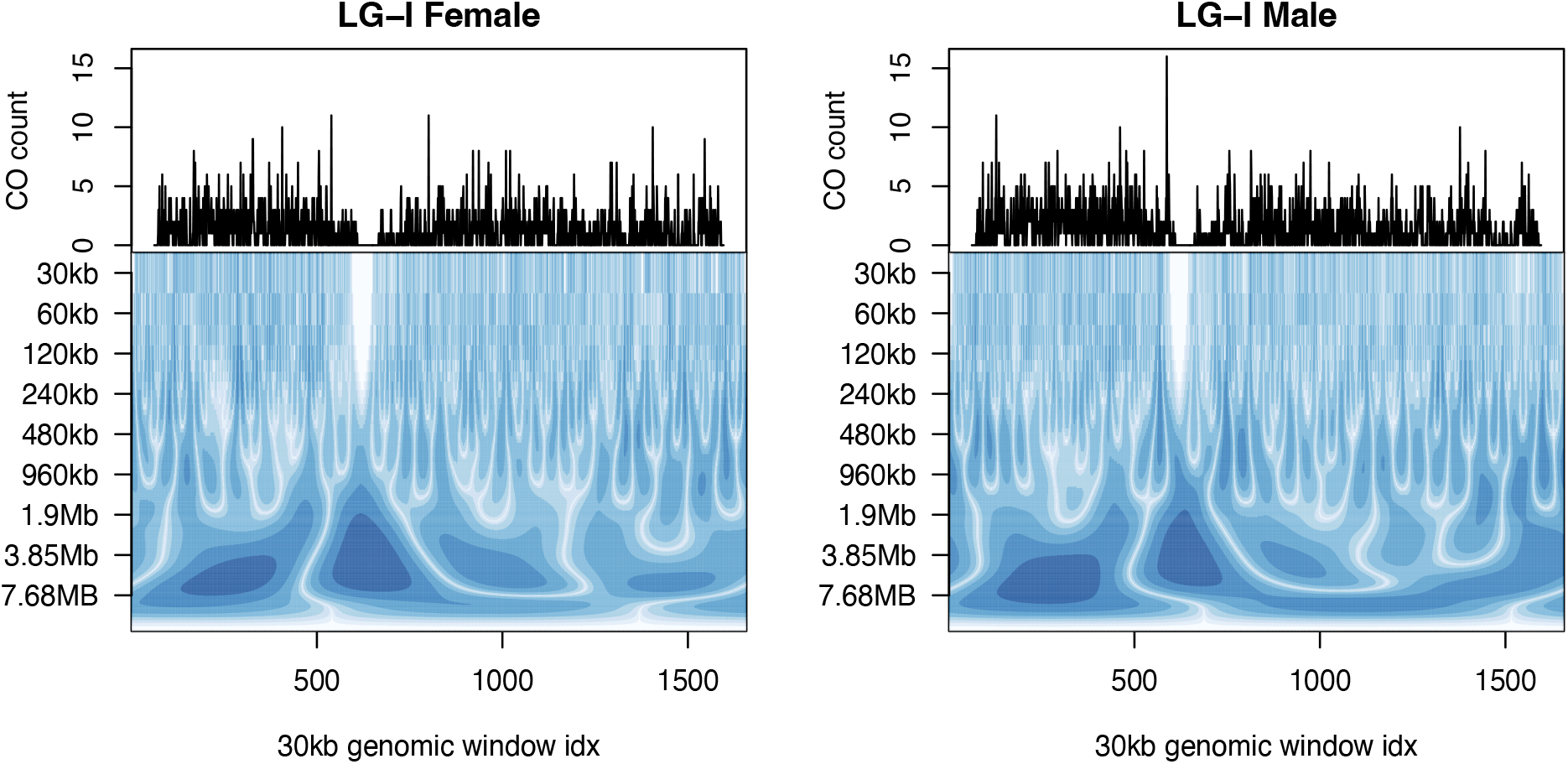
Continuous wavelet transform (CWT) of female and male cumulative CO count for LG-I. Line graphs display cumulative CO counts for each sex (in columns) across 30 kb genomic windows. Wavelet coefficients at each scale using a CWT is displayed as a power spectrum at each scale.

### Pairwise LD vs. cumulative CO count

In order to evaluate the legitimacy of using CO counts as a proxy for recombination rate, cumulative CO count within 960-kb non-overlapping genomic windows was compared to mean pairwise LD within the same set of windows in a subpopulation of 220 trees collected from the same region as the parents of the 7×7 cross (Figure S1). As expected, a strong negative correlation was observed between cumulative CO counts and mean pairwise LD (*r*_spearman_ = −0.69; p = 4.0×10^-4^), consistently across all of the 19 linkage groups (Figure S9).

### Prediction of localized CO pattern using genomic correlates

As is the general trend for plants, CO counts were lower in proximity to inferred centromere and telomere positions and higher at pericentromeric regions when chromosomes were metacentric or submetacentric. Genomic features such as repeat content (*Gypsy-*LTR, *Copia-*LTR or simple repeats) GC-content and gene content were significantly associated with expected cumulative CO counts in 960-kb intervals (Model-1 cross-validation R^2^ = 0.52, RMSE = 12.93). *Gypsy-*LTR, percent-GC and *Copia-LTR* content all had significant negative effects on cumulative CO counts while gene content and simple repeats were positively associated with CO counts (Table S9). Dropping simple repeat content from the linear model increased the magnitude of the effects for other factors which may be explained by the multicollinearity exhibited between these variables (Figure S10). Dropping simple repeats from the prediction model also had minimum effect of the predictability of local CO counts within these genomic windows (Model-2 cross-validation R^2^ = 0.52, RMSE = 13.03).

### Fine-scale differences in CO counts between sexes

Chromosome-scale differences in CO counts between sexes (**Figure 3a**), were projected to a finer scale within chromosomes. Informed by the wavelet analysis, a non-overlapping window size of 960 kbps was used in the fine-scale analysis. Differences in cumulative CO counts between males and females showed similar trends across the genome (*r*^2^_Spearman_ = 0.78; p-value < 2.2×10^-16^) and were not qualitatively different (Kolmogorov-Smirnov test, p = 0.6093, **Figure 2**, Figure S6). Nevertheless, sixteen genomic windows were identified (Exact-Poisson test; FDR_BH_ ≤ 0.25) that show differences in CO counts between females and males, spread through the genome (**Figure 3b**). Interestingly, only one genomic window showed higher CO counts for females (**Figure 3b**). The overwhelming majority of windows showed higher CO rates for males. Furthermore, there were long consecutive chromosome segments for which male CO counts were consistently higher than females, such as at the beginning of LG-1, though the differences were not statistically significant for individual intervals (**Figure 3b**). Windows with a higher number of COs in males had lower GC content (Wilcoxon Rank-sum test; W = 655; p = 0.057) and gene content (Wilcoxon Rank-sum test; W = 623; p = 0.034) compared to windows that did not show significant sex bias. None of the other variables related to repeat content (*Copia, Gypsy* or simple repeats counts) had any significant difference (FPR ≤ 0.05) between these two groups (**Figure 4**).

**Figure 3:**
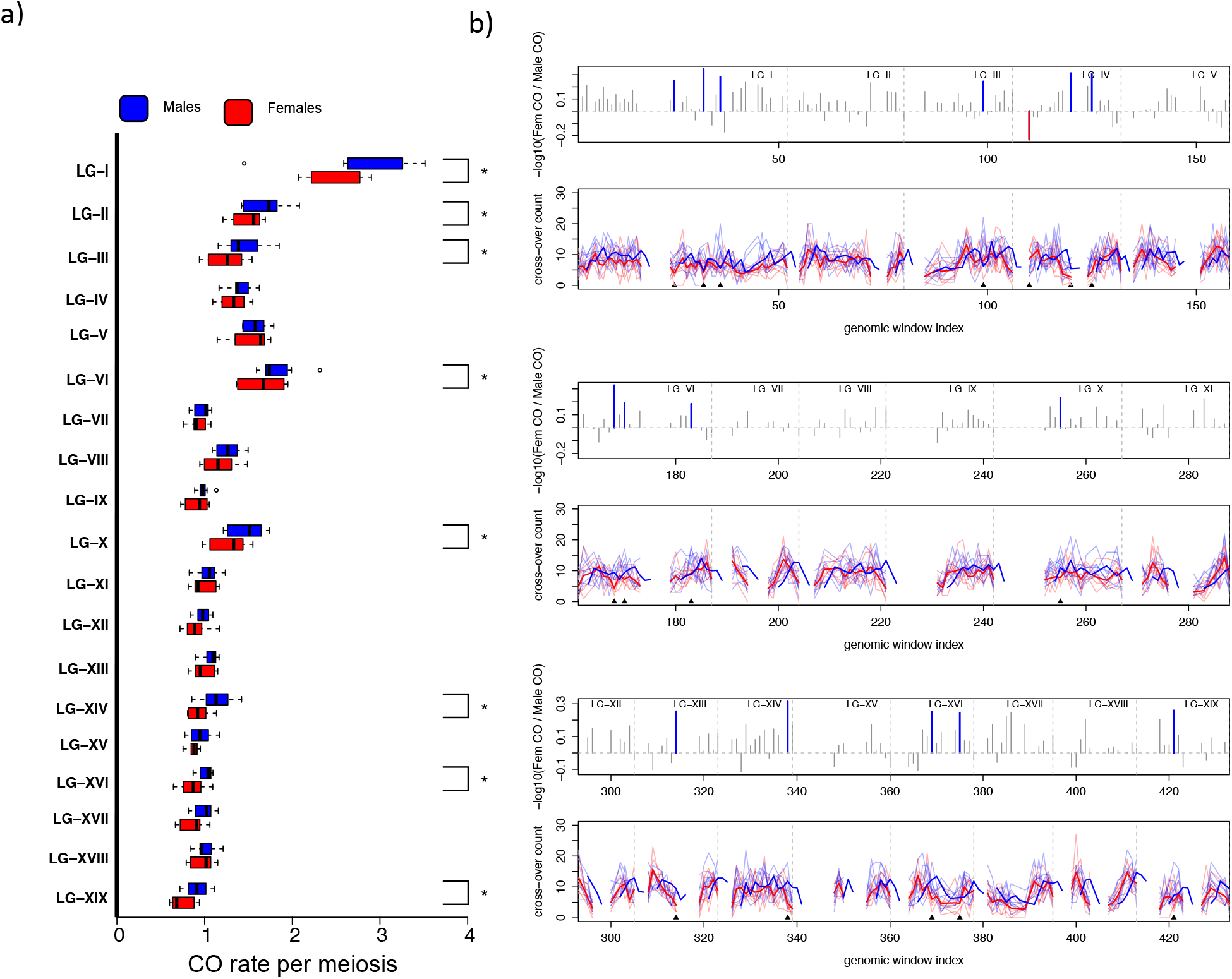
Broad to fine scale CO count differences between sexes: **a)** CO rate per chromosome per meiosis in males (blue) vs. females (red). LGs marked with an asterisk shows significant differences in cumulative CO counts between the sexes (χ^2^_df=1_; FWER ≤ 0.05). **(b)** Fine scale differences in recombination between sexes are shown. Log_10_ transformed male to female cumulative CO count ratios are shown in grey bars for each window and blue and red bars indicate male biased and female biased heterochiasmy respectively. Only one window showed female bias. Mean CO count shown for male (thick blue line) and female (thick red line) parents. Lighter colored lines represent counts for individual parents of each sex. Gaps indicate centromeric or telomeric regions that were removed from our statistical analysis due to sparsity of counts.

**Figure 4:**
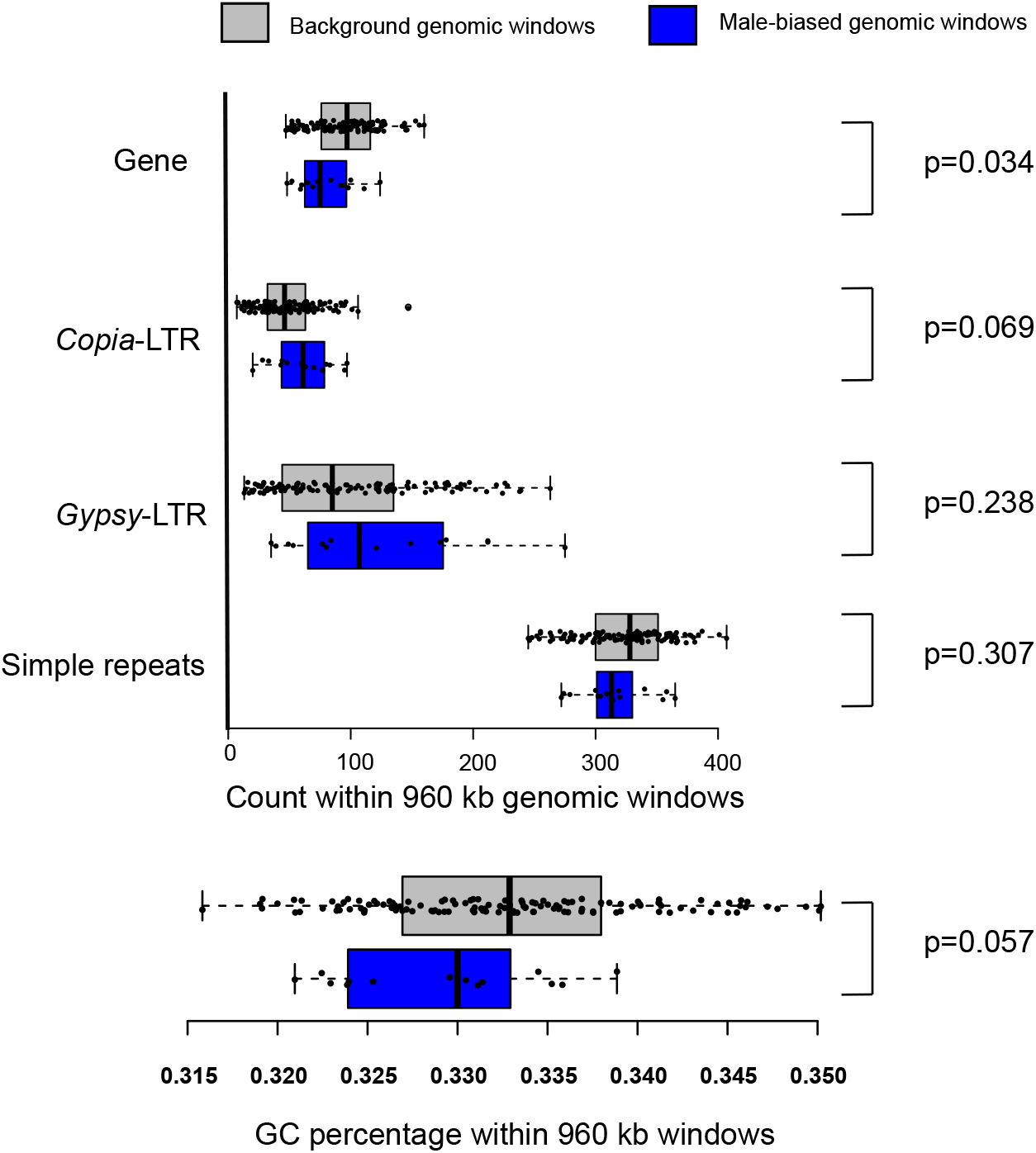
Genomic features associated with elevated CO counts in males: Enrichment of genomic features in windows that showed significantly elevated CO counts were compared with windows with background male/female ratios.

### Identification of genome-wide CO hotspots

At yet a finer resolution of 30 kb, the observed distribution of CO counts showed an excess of windows with zero CO events, most likely due to windows that overlap centromeres and a longer than expected tail that may consist of candidate CO hotspots (Figure S11). CO hotspot regions were defined as 30-kb windows that showed significantly elevated cumulative CO counts in comparison to the genome-wide average. Initially 119 candidate CO hotspot regions were identified (λ = 2.98, FWER_Bonferroni_ ≤ 0.05), which were short-listed to 67 windows by applying minimum thresholds for flanking number of markers and index-of-dispersion across half-sib families for CO counts in order to enrich for true-positives. These robust CO hotspot candidates were spread across the genome in clusters of adjacent genomic regions (**Figure 5a**). Genome-wide hotspots were evaluated separately for males and females with adjusted CO-counts per window (λ_female_ = 1.38; λ_male_ = 1.54) where 38 and 24 CO hotspot windows were identified respectively (FWER_Bonferroni_ ≤ 0.05). Of these, 21 and 11 windows overlapped with hotspots identified with females and males combined, respectively. Although female and male hotspots were within the general vicinity of each other, only a single CO hotspot window was shared between the sexes in our analysis (Figure S12). However, the lower number of CO hotspot windows detected for males in comparison to females could be due to the higher null expectation and variance of CO counts for males leading to a more stringent FWER threshold for meeting statistical significance in comparison to females.

**Figure 5:**
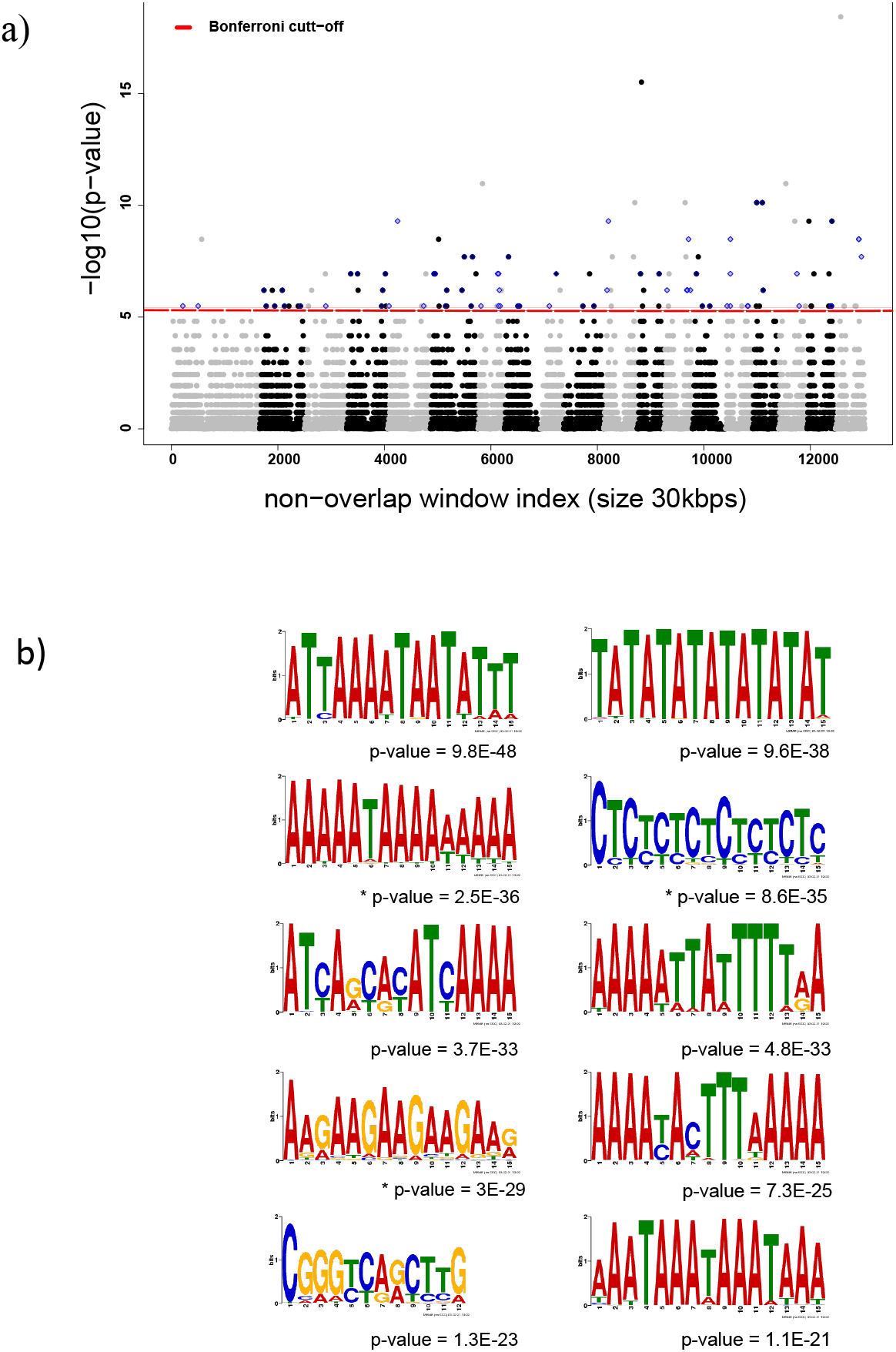
Candidate cross-over (CO) hotspot regions. **(a)** Points marked as blue diamonds represent non-overlapping windows of size 30kbps that represent cross-over counts (CO) hotspots based on a probability model that describes genome-wide average CO per window by a Poisson null distribution (λ=3; FWER ≤ 0.05). **(b)** The top ten DNA sequence motifs enriched in CO regions that were demarcated to a genomic region of 10 kbps or less. P-values denoted with a ‘*’ identifies DNA sequence motifs that have significant blast hits to recombination hotspot associated DNA sequence motifs previously identified for Arabidopsis and wheat.

The enrichment of genomic features in CO hotspots was evaluated through comparison with windows with background levels of CO counts using a two tailed Wilcoxon rank sum test (Figure S13). Simple repeat count was most strongly elevated in hotspots (Wilcoxon Rank-sum test; W = 67698; p = 1.795 x 10^-7^), while GC content (Wilcoxon Rank-sum test; W = 130366; p = 0.003048) and *Copia*-LTR content (Wilcoxon Rank-sum test; W = 124020; p = 0.0198) were significantly lower. Neither *Gypsy-*LTR nor gene content showed significant differences (FPR ≥ 0.05).

### DNA sequence motifs associated with COs

In order to investigate whether there are DNA sequence motifs associated with higher CO probability, the analysis was restricted to narrowly delimited CO regions. A total of 199 DNA sequence motifs putatively associated with COs are reported and of those, motifs that ranked within the first three were all A/T-rich repeat sequences (**Figure 5b**). Interestingly, most of the enriched DNA sequence motifs identified within the top 10 showed considerable abundance across the whole genome. The full list of enriched DNA sequence motifs and matching or similar sequences across the genome is detailed in Table S10. The enriched DNA sequence motifs also showed high similarity to such motifs associated with recombination hotspots in wheat and *Arabidopsis* as well (**Figure 5b**; Table S11 & Table S12). However, our analysis was not able to identify any DNA sequence motifs that are associated with CO hotspots when compared to regions with average COs at a maximum resolution of 30 kb.

## DISCUSSSION

### Fourteen high-fidelity genetic maps

We reported 14 dense framework genetic maps each with 19 LGs, corresponding to the accurate haploid number of chromosomes in *P. trichocarpa*, all of which were derived from intra-specific crosses. Full-sib family sizes were too small for accurate genetic mapping, so we used half-sib families for map construction (Grattapaglia *et al*. 1996; Littrell *et al*. 2018). Given that each parent was crossed with the same set of individuals from the opposite sex in a full-factorial cross, genetic maps among individuals within a sex are highly comparable with each other. The concordance, collinearity, and the congruity of our genetic maps can be mainly attributed to key steps in our pipeline such as 1) pedigree error correction 2) robust variant calling pipeline 3) genotype error correction and imputation of parents 4) phasing and identification of haplotype configuration in offspring and 5) constructing the initial framework maps using a moderate number of intensively filtered markers to avoid map inflation. Although whole-genome resequencing provided a very high density of SNP and INDEL markers distributed across the genome, the resolution of our maps was largely limited by the number of meioses within each half-sib family, which resulted in a large number of markers being binned due to redundant genetic information (Da *et al*. 1998).

We reported comparable recombination rates across LGs and considerable variation in recombination rate among individuals. Our approach simulates a structured tree breeding program and has high probability of identifying large haplotype blocks with a history of limited internal recombination (Jansen *et al*. (2003); Di Pierro *et al*. (2016). Such information is potentially useful to breeding programs because quantitative trait loci (QTL) falling within these regions can be targets for marker assisted selection (Wu *et al*. 1998; Muranty *et al*. 2014). Fine-scale estimates of recombination rates and exact recombination breakpoints identified in this study will be useful in QTL delineation for feedstock relevant phenotypes. These estimates provide an important foundation for accelerated domestication as well as basic investigations into the evolutionary biology and molecular genetics of this model forest tree (Grattapaglia and Resende 2011; Isik 2014).

### Higher rates of CO in males

One of our most striking findings was the higher rates of CO in male parents of *P. trichocarpa* (male-biased heterochiasmy) where we had an equal number of gametes (829) for each sex. A higher rate of recombination in males had been previously reported for an interspecific cross of *Populus* (Yin *et al*. 2002). However, a revisit of these data in a metanalysis failed to confirm the finding (Lenormand and Dutheil 2005). A possible reason could be that the metanalysis only considered markers that were shared between the male and female maps, which also resulted in overall shorter cumulative map lengths. In our study, we compared cumulative CO counts in each LG accounting for individual variance and other covariates such as LG identity and half-sib family size, after which 8 out of 19 LGs showed male-biased heterochiasmy at the chromosomal scale.

We also observed higher CO counts in male gametogenesis in an overwhelming majority of the 960-kb windows analyzed genome-wide. Overall, male and female CO landscapes were highly correlated for a given LG and were not qualitatively different, unlike in *Arabidopsis* (Lloyd and Jenczewski 2019), human (Kong *et al*. 2010) or mouse (Brick *et al*. 2018) where patterns of CO distribution are dissimilar between sexes. This suggests a model where the CO landscape is spatially conserved across sexes in *P. trichocarpa*, at least at 960-kb resolution. Heterochiasmy is commonly observed across a wide range of organisms, including plants (Lenormand and Dutheil 2005; Sardell and Kirkpatrick 2020). It is a fast-evolving trait, as evidenced by the observation that phylogenetic inertia does not limit its emergence (Lenormand and Dutheil 2005). Five possible explanations have been proposed for the observed differences in CO counts between sexes: **1)** cellular, molecular mechanisms; 2) pleiotropic effects of the sex chromosome or Sex Determination Region (SDR); **3)** sex dimorphism in haploid selection (Hunt and Hassold 2002; Petkolv *et al*. 2007; Lloyd and Jenczewski 2019); **4)** meiotic drive mechanisms (Brandvain and Coop 2012); and **5)** differential external factors during meiosis (Phillips *et al*. 2015; Coulton *et al*. 2020);. Evidence for each of these is discussed below.

Female and male meioses are fundamentally different in plants and could be driven by cellular and molecular processes that maintain a differential advantage in sexes. Heterochiasmy can result from differences in the amount and distribution of DSBs, their resolution and maturation as either CO or NCO as well as the sensitivity to CO interference (Capilla-Perez *et al*. 2021). The length of the synaptonemal complex (SC), which is a structure formed during meiosis and responsible for ensuring proper chiasma formation and maintenance, is shown to have a positive correlation to CO occurrence in *Arabidopsis* (Drouaud *et al*. 2007; Giraut *et al*. 2011), leading to higher rates of recombination in male gametogenesis. Furthermore, SCs are deemed essential in Arabidopsis for CO interference, and CO frequencies are shown to equalize in the absence of SCs (Capilla-Perez *et al*. 2021). Heterochiasmy in *Zea mays* also reveals more COs in male gametogenesis and more hotspots. This varies by genotype and has been attributed to variation in CO maturation, which could also be related to differences in synaptonemal complex length (Luo *et al*. 2019). Differential methylation and chromatin structure have also been implicated for sex differences in recombination (Wang and Copenhaver 2018).

In *Populus balsamifera* which is a sister species to *P. trichocarpa* (Rood *et al*. 1986; Eckenwalder 1996; Wang *et al*. 2020) male biased expression in a subset of chromatin remodeling and DNA methylation regulator genes have been observed (Cronk *et al*. 2020). Although we did not directly investigate chromatin modifications or DNA methylation in our study, we reported that observed male-biased heterochiasmy had close association with gene content and percent GC content in 960-kb windows. These factors could themselves determine specific locales and amount of DNA methylation or chromatin modifications shaping the quantitative differences we observed. However, this remains an open question, and the underlying molecular mechanisms that are responsible for these differences have yet to be identified (Bergero *et al*. 2021).

The *Populus* sex determination region (SDR) contains at least two genes involved in differential DNA methylation between sexes (Geraldes *et al*. 2015; Zhou *et al*. 2020; Kim *et al*. 2021). However, whether these genes could have pleiotropic effects that extend to other regions of the genome to affect DNA methylation and histone remodeling is still unclear (Sardell and Kirkpatrick 2020). Intriguingly, a study of DNA methylation across a range of tissues in *P. trichocarpa* found the highest rates of methylation in male floral tissue (Vining *et al*. 2012), though the significance of this for chromatin structure and CO occurrence is unclear. On the other hand, not all chromosomes or fine-scale genomic-windows in our study showed male biased heterochiasmy, which means that the pleiotropy-based explanation does not completely address our observations for windows in which female rates were higher or in which no significant bias was observed either way.

Another potential contributor to heterochiasmy in plants is differential opportunity for haploid selection (Lenormand and Dutheil 2005). In plants there is a substantial time period in which haploid spores, unmasked from effects of homologous partners, may express their genes that are then subject to selective pressures. In anemophilous species such as *P. trichocarpa* the opportunity for haploid selection and the distance of dispersal may favor mechanisms that promote genetic variation and thus increased shuffling of haploid genomes (Slavov *et al*. 2009). Since male gametes typically are dispersed much further than female gametes in trees, including in *Populus* (Slavov *et al*. 2009; Slavov *et al*. 2012), this mechanism could account in part for the observed male-biased heterochiasmy. However, increased recombination may not be favored when recombination breaks-up favorable epistatic interactions thus reducing the mean fitness of the resulting population in subsequent generations (Charlesworth and Barton 1996). In angiosperms, males are subjected to increased selective pressure due to pollen competition and should in theory have lower recombination rates that preserve favorable haplotype combinations minimizing recombination load (Lenormand and Dutheil 2005). However, the validity of this theory may not apply to *Populus*, which has several potentially mitigating characteristics, including dioecy, anemophily, and extensive clonal reproduction through sprouting and rooting of woody propagules.

Recombination rate modifiers that alter the efficiency of sex specific meiotic drive mechanisms have also been implicated for sex-based differences in recombination rate. Although specific mechanistic processes are still unclear, a leading hypothesis based on population genetic modelling suggests that female meiotic drive could either enhance or suppress recombination rates, based on the interaction of whether the drive system functions during meiosis-I (MI) or meiosis-II (MII) and whether it is linked or unlinked to a recombination rate modifier (Brandvain and Coop 2012). As per this reasoning, relatively higher rates of recombination in males as we observed could be the result of an active suppression of recombination in females due to either an MI meiotic drive system in phase with a recombination suppressor or an MII drive system unlinked to a recombination suppressor during female gametogenesis. However, under this model, recombination rate differences between males and females should be more prominent closer to the centromeric region which was not observed in our study. Although an MII meiotic driver system (Ab10) affecting recombination has been characterized in maize (Hiatt and Dawe 2003), evidence for such systems are not reported for any *Populus* species. Female biased sex ratios are observed in a number of Salicaceae species, and even here the only mechanism identified does not implicate sex chromosome meiotic drive (Pucholt *et al*. 2017).

Higher recombination rates for male gametogenesis have been reported in interspecific crosses of *Populus* (Yin et al. 2002), as well as within male strobili of *Pinus* (Groover *et al*. 1995; Plomion and Omalley 1996), and in the latter it may be attributed to the temporal differences in meiosis between male and female strobili where external temperature differences during gametogenesis are driving sex differences (Moran *et al*. 1983). Details or direct evidence of differential external or internal environmental effects during gametogenesis are not reported for *P. trichocarpa* other than possible ambient chemical composition differences within catkins.

### Genome-wide cross-over hotspots

Conserved DNA sequence motifs associated with CO hotspots have been identified in domesticated plants such as *Zea mays*, *Triticum aestivum* (Darrier *et al*. 2017), and *Oryza sativa* (Mezard 2006), yet similar studies on undomesticated forest trees are rare (Slavov *et al*. 2012; Silva - Junior and Grattapaglia 2015; Apuli *et al*. 2020). Although LD-based recombination rate estimates have been previously obtained for *P. trichocarpa* (Slavov *et al*. 2012), these estimates are subject to stochasticity of population demography and do not provide the flexibility to decompose sex-based differences. This study revealed the genome-wide recombination landscape and patterns of inheritance at a finer scale than has previously been possible in a single generation of *Populus*, and rarely achieved in undomesticated plants. Using high density phased and imputed SNPs, we were able to delineate recombination breakpoints within 30-kb windows. We identified 67 windows enriched for recombination hotspots and these sites displayed a genome-wide distribution yet clustered away from putative centromeres and telomeres. A role for absence of chromatin compaction and trimethylation of lysine 4 in histone H3 has been identified as strong markers of DSBs and recombination in yeast and mammals (Pan *et al*. 2011; Smagulova *et al*. 2011; Lange *et al*. 2016). However, such patterns cannot be generalized in plants such as *Zea mays* in which association between DSBs and recombination has been shown to be moderated by other factors such as the amount of repetitive DNA (Rodgers-Melnick *et al*. 2016; He *et al*. 2017). Molecular mechanisms that regulate DSBs and subsequent recombination are complex and involve the interaction of localized genomic features related to chromatin compaction such as histone remodeling, DNA-methylation, gene content, and repeat content in *Arabidopsis* (Yelina *et al*. 2012; Shilo *et al*. 2015).

Regions with high AT richness have been associated with elevated CO frequency in *A. thaliana* (Choi *et al*. 2018). Poly(dA:dT) tracts and their flanking regions are shown to be depleted of nucleosomes *in-vitro*, in addition to blocking the spread of post-translational histone modifications that limit accessibility to recombination and transcriptional machinery (Segal and Widom 2009). Similarly, hotspots identified in our study were also associated with higher AT content and were comparatively enriched for simple repeat elements. This observation is further supported by the association of AT-rich DNA sequence motifs at CO sites. Although not exact matches, we reported several DNA sequences that showed close similarity to such sequences that were observed in wheat (Darrier *et al*. 2017) and Arabidopsis (Shilo *et al*. 2015) that are associated with DNA methylation and histone remodeling.

We also presented a statistical model that accounts for 53% of the variation in CO counts in 960-kb non-overlapping windows using localized genomic correlates. Consistent with other plant species (Mezard 2006; Gaut *et al*. 2007), we observed that COs tend to be higher in regions enriched for genes and simple repeats while negatively influenced by higher percent GC content along with LTR-*Gypsy* and LTR-*Copia*. It is difficult to parse out the multicollinearity among these explanatory variables and by excluding the centromeric and telomeric regions we have tried to minimize this issue. Phylogenetically independent contrasts also show similar trends as we observed (Tiley and Burleigh 2015) and suggest fundamental genetic or evolutionary driving forces acting on CO control. It is hypothesized that transposable elements (TEs) accumulate within regions with low recombination rates, which initiates a positive feedback loop spreading recombination suppression to adjacent regions in a model that describes coevolutionary interaction (Kent *et al*. 2017). We observed a positive correlation between gene density and CO counts that concur with general tendency in plant species which provides a stark contrast to the absence of a conclusive relationship observed in animals and fungi at these broader scales (Mezard 2006; Gaut *et al*. 2007; Haenel *et al*. 2018). This is in line with the recombination modifier theory for the existence of CO hotspots where mutations of sequence variants that promote shuffling of flanking regions are favored, since such a situation makes selection and local adaptation more efficient especially in the case of perennial forest trees (Hey 2004).

## CONCLUSIONS

Our analysis revealed the genome-wide recombination landscape at a finer scale than has previously been possible in most undomesticated woody plant species. In *P. trichocarpa*, this information will be used to delineate Quantitative Trait Loci (QTL) for phenotypes of breeding relevance (height, diameter, bud set, and disease resistance) and along with recombination rate estimates to improve genomic prediction models. Given that *P. trichocarpa* is a promising renewable feedstock for bioenergy and bioproducts, we believe that our findings on recombination and CO rate estimates will be useful for ongoing efforts to accelerate domestication of this and other feedstocks, as well as future studies that investigate broader questions such as evolutionary history, perennial development related to phenology, wood formation, vegetative propagation, and dioecy that cannot be studied using conventional plant model systems such as *Arabidopsis*, rice, or maize.

## AUTHOR CONTRIBUTIONS

CRA carried out genetic mapping and subsequent data analysis and wrote the manuscript. DMS provided advice on genetic mapping, data analyses, performed data analyses, wrote and edited the manuscript. RZ provided methods and scripts for estimating repeats content, gene content and AT/GC composition within genomic windows. KB led the resequencing of the genomes of individuals in this study at JGI. CD, MZ, YY and AL carried out DNA sequence library preparation, sequencing and quality control. BS designed the 7×7 cross. KH generated the 7×7 crosses and managed the clonal progeny field trial. GAT provided funding and leadership for the project. SPD led the study and helped write and edit the manuscript. All authors read and approved the final manuscript.

## ACKNOWLEDGMENTS

The work (proposal: 10.46936/10.25585/60001012) conducted by the U.S. Department of Energy Joint Genome Institute (https://ror.org/04xm1d337), a DOE Office of Science User Facility, is supported by the Office of Science of the U.S. Department of Energy operated under Contract No. DE-AC02-05CH11231.The authors would like to acknowledge the High-Performance-Computing group at West Virginia University, funded in part by the National Science Foundation EPSCoR Research Infrastructure Improvement Cooperative Agreement #1003907, the state of West Virginia (WVEPSCoR via the Higher Education Policy Commission) and WVU.

Finally, we thank researchers from the Center for Bioenergy Innovation who contributed to this work.

## DATA AVAILABILITY STATEMENT

Raw sequence reads are publicly available at the JGI genome portal under (proposal: 10.46936/10.25585/60001012) and metadata related to sequence read archive (SRA) accessions used in this study are provided in Table S13.

## SUPPLEMENTARY INFORMATION

**Supplementary methods:** Discrete wavelet analysis for identifying the scales at which sex-based CO count differences are most prominent

**Table S1:** Design of crosses used for genetic map construction

**Table S2:** Criteria for hard-filtering variants

**Table S3:** Evaluation of statistical models for expected count of COs

**Table S4:** Statistical models for predicting localized CO counts

**Table S5:** Genetic linkage maps for all 14 half-sib families

**Table S6:** Summary statistics for median female and male genetic maps

**Table S7**: Chi-square test for equal CO counts between females and males

**Table S8:** Statistical model parameter estimates

**Table S9:** Linear model for broad-scale genomic features (in non-overlapping windows of 960 kbps) that explains cumulative CO counts

**Table S10:** Enriched DNA sequence motifs in CO regions and their abundance in the genome

**Table S11:** DNA sequence motifs associated with recombination hotspots in prevous publications

**Table S12:** Similarity of DNA sequence motifs identified in our study to such sequences associated with recombination hotspots as reported in prevous publications

**Table S13:** Sequence reads metadata

**Figure S1:** Geographical location of parental clones in the 7×7 cross.: The mapping population we use here is a subset of the GWAS population (Wegrzyn *et al*. 2010) in western USA and Canada which include individuals from the whole natural range Washington, Oregon and California in The USA and British Columbia, Canada. Circles indicate where the GWAS clones were collected and the 14 red circles indicate parents of the 7×7 trial. The yellow star marks the location of the *Populus trichocarpa* male collected from the Mount Hood region used as the reference genome (Stettler-14) in this study.

**Figure S2:** Haplotype phasing and imputation. a) Observed and b) imputed haplotypes, for offspring of the half-sib family of female parent GW-1863 on LG-III. Offspring were phased using trio genotype information. red or blue lines represent each of the maternal haplotypes. Missing genotypes and Mendelian violations are marked in yellow. This process was completed for all 19 LGs for each of the 14 parents.

**Figure S3:** (a) CO region size distribution for all 19 LGs for all half-sib families; (b) median physical distance between neighboring markers in all genetic maps; (c) median genetic map distance between neighboring markers for all genetic maps.

**Figure S4:** Statistical model fit for GLMM analyzing chromosome scale sex-based differences in CO counts: The distribution of Pearson residuals derived after fitting the GLMM with a log-link function did not show any significant trends across fitted values (a) or any of the explanatory variables (b & e) such as half-sib family identity, LG identity, sex or physical length of the chromosome

**Figure S5:** Partitioned wavelet variance. Histograms (grey shading) represent null distribution of partitioned wavelet variance at each scale from 60 kb through 3.85 Mb (in rows), for four selected chromosomes (in columns). Null distributions were derived by randomly assigning parents to nominal groups of sex while preserving spatial structure of windows within each chromosome. The red and blue vertical lines represent the observed wavelet variance at each scale for female and male groups respectively. Significant separation between red and blue lines from 60 kb through 3.85 Mb are characteristic of LGs in which heterochiasmy is present such as LG-I, LG-XIV, while others such as LG-XI and LG-XIII show no such trend and do not show differences between sexes.

**Figure S6:** Power spectrum of continuous wavelet transformation (CWT). LG-II through LG-XIX (rows) are shown for male and female (columns) cumulative CO counts at each scale ranging from 30 kb to 7.68 MB.

**Figure S7:** A high correlation was observed between genetic map averages produced in this study in comparison to similar maps from an independent study using SSR markers (Yin *et al*. 2004). Both male averaged (blue) and female averaged (red) map sizes showed high correlation (median female maps: r = 0.92, p= 1.709 x 10^-8^; median male maps: 0.93, p= 5.868 x 10^-9^). Blue and red dashed lines represent the linear regression fit for male and female median genetic maps for 19 linkage groups respectively.

**Figure S8:** Regression of mean CO count vs. physical length for a given chromosome. Dots represent the mean CO count for a given half-sib family for a given chromosome. The dashed blue line represents the linear regression fit. Physical length of the chromosome explained majority of the variation in mean CO counts (R2 = 0.77, p=2.2 x10^-16^). There is considerable variance between half-sib families for a given chromosome and the magnitude increases as the size of the chromosome increases.

**Figure S9:** Comparison of pairwise LD and cumulative CO counts: A comparison of pairwise LD and cumulative CO counts across genomic windows of size 960 kb. A strong negative correlation was observed between cumulative CO counts and mean pairwise LD (r_spearman_= −0.711, p=2.2 x 10^-16^).

**Figure S10:** CO pattern prediction using genomic correlates at 1Mb scale. This shows the pair-wise relationship between various explanatory genomic features used to predict CO count within a given genomic region of size 1Mbps. There is high correlation among LTR families, as well as among LTRs, gene content, and GC content. High multi-collinearity is expected when these variables are used in linear models.

**Figure S11:** Empirical (blue) vs. null (orange) distributions of CO counts per 30 kb non-overlapping window size across the genome. The null distribution assumes that COs are distributed independently and identically across the genome, conforming to a theoretical Poisson distribution (λ_cumulative_= 2.98). Observed CO counts per window are enriched for zero counts most likely due to windows that overlap centromeric span while longer than expected tail region indicate CO hotspot candidates

**Figure S12:** CO hotspots for males and females. Genome-wide hotspots were evaluated separately for males (blue) and females (red) with adjusted CO-counts per window for males and females and 38 and 24 CO hotspot windows were identified respectively (λ_female_ =1.38; λ_male_ =1.54; FWER_Bonferroni_ ≤ 0.05). 21 and 11 windows overlapped with genomic scale cumulative CO-count hotspots for females and males respectively. Female and male hotspots were within the general vicinity of each other.

**Figure S13:** Genomic correlates that influence localized CO patterns. Genomic correlates associated with CO hotspots, in 30 kb genomic windows that showed significantly elevated CO counts (hotspots) with windows with background levels of CO counts.

## REFERENCES

Altschul, S. F., W. Gish, W. Miller, E. W. Myers and D. J. Lipman, 1990 Basic local alignment search tool. Journal of Molecular Biology 215: 403–410.

Apuli, R. P., C. Bernhardsson, B. Schiffthaler, K. M. Robinson, S. Jansson et al., 2020 Inferring the genomic landscape of recombination rate variation in European Aspen (*Populus tremula*). G3-Genes Genomes Genetics 10: 299–309.

Auton, A., and G. McVean, 2007 Recombination rate estimation in the presence of hotspots. Genome Research 17: 1219–1227.

Bailey, T. L., M. Boden, F. A. Buske, M. Frith, C. E. Grant et al., 2009 MEME SUITE: tools for motif discovery and searching. Nucleic Acids Research 37: W202–W208.

Baudat, F., J. Buard, C. Grey, A. Fledel-Alon, C. Ober et al., 2010 PRDM9 is a major determinant of meiotic recombination hotspots in humans and mice. Science 327: 836–840.

Bauer, E., M. Falque, H. Walter, C. Bauland, C. Camisan et al., 2013 Intraspecific variation of recombination rate in maize. Genome Biology 14: 17.

Benjamini, Y., and Y. Hochberg, 1995 Controlling the false discovery rate - a practical and powerful approach to multiple testing. Journal of the Royal Statistical Society Series B-Statistical Methodology 57: 289–300.

Bergero, R., P. Ellis, W. Haerty, L. Larcombe, I. Macaulay et al., 2021 Meiosis and beyond - understanding the mechanistic and evolutionary processes shaping the germline genome. Biological Reviews 96: 822–841.

Bherer, C., C. L. Campbell and A. Auton, 2017 Refined genetic maps reveal sexual dimorphism in human meiotic recombination at multiple scales. Nature Communications 8: 9.

Bradshaw, H. D., R. Ceulemans, J. Davis and R. Stettler, 2000 Emerging model systems in plant biology: Poplar (*Populus*) as a model forest tree. Journal of Plant Growth Regulation 19: 306–313.

Bradshaw, H. D., M. Villar, B. D. Watson, K. G. Otto, S. Stewart et al., 1994 Molecular genetics of growth and development in Populus. III. A genetic linkage map of a hybrid poplar composed of RFLP, STS, and RAPD markers. Theoretical and Applied Genetics 89: 167–178.

Brandvain, Y., and G. Coop, 2012 Scrambling eggs: meiotic drive and the evolution of female recombination rates. Genetics 190: 709–723.

Brick, K., S. Thibault-Sennett, F. Smagulova, K. W. G. Lam, Y. M. Pu et al., 2018 Extensive sex differences at the initiation of genetic recombination. Nature 561: 338–342.

Burt, A., G. Bell and P. H. Harvey, 1991 Sex-differences in recombination. Journal of Evolutionary Biology 4: 259–277.

Capilla-Perez, L., S. Durand, A. Hurel, Q. C. Lian, A. Chambon et al., 2021 The synaptonemal complex imposes crossover interference and heterochiasmy in *Arabidopsis*. Proceedings of the National Academy of Sciences of the United States of America 118: 11.

Charlesworth, B., and N. H. Barton, 1996 Recombination load associated with selection for increased recombination. Genetical Research 67: 27–41.

Charlesworth, D., B. Charlesworth and G. Marais, 2005 Steps in the evolution of heteromorphic sex chromosomes. Heredity 95: 118–128.

Chhetri, H. B., D. Macaya-Sanz, D. Kainer, A. K. Biswal, L. M. Evans et al., 2019 Multitrait genome-wide association analysis of *Populus trichocarpa* identifies key polymorphisms controlling morphological and physiological traits. New Phytologist 223: 293–309.

Choi, K., and I. R. Henderson, 2015 Meiotic recombination hotspots - a comparative view. Plant Journal 83: 52–61.

Choi, K., X. H. Zhao, A. J. Tock, C. Lambing, C. J. Underwood et al., 2018 Nucleosomes and DNA methylation shape meiotic DSB frequency in Arabidopsis thaliana transposons and gene regulatory regions. Genome Research 28: 532–546.

Coulton, A., A. J. Burridge and K. J. Edwards, 2020 Examining the effects of temperature on recombination in wheat. Frontiers in Plant Science 11: 1–13.

Cronk, Q., R. Soolanayakanahally and K. Brautigam, 2020 Gene expression trajectories during male and female reproductive development in balsam poplar (*Populus balsamifera* L.). Scientific Reports 10: 14.

Da, Y., P. M. VanRaden, N. Li, C. W. Beattie, C. X. Wu et al., 1998 Designs of reference families for the construction of genetic linkage maps. Animal Biotechnology 9: 205–228.

Dapper, A. L., and B. A. Payseur, 2017 Connecting theory and data to understand recombination rate evolution. Philosophical Transactions of the Royal Society B-Biological Sciences 372: 11.

Darrier, B., H. Rimbert, F. Balfourier, L. Pingault, A. A. Josselin et al., 2017 High-resolution mapping of crossover events in the hexaploid wheat genome suggests a universal recombination mechanism. Genetics 206: 1373–1388.

DePristo, M. A., E. Banks, R. Poplin, K. V. Garimella, J. R. Maguire et al., 2011 A framework for variation discovery and genotyping using next-generation DNA sequencing data. Nature Genetics 43: 491–498.

Di Pierro, E. A., L. Gianfranceschi, M. Di Guardo, H. J. J. Koehorst-van Putten, J. W. Kruisselbrink et al., 2016 A high-density, multi-parental SNP genetic map on apple validates a new mapping approach for outcrossing species. Horticulture Research 3: 13.

Drouaud, J., R. Mercier, L. Chelysheva, A. Berard, M. Falque et al., 2007 Sex-specific crossover distributions and variations in interference level along *Arabidopsis thaliana* chromosome 4. Plos Genetics 3: 1096–1107.

Eckenwalder, J. E., 1996 Systematics and evolution of Populus, pp. 7–30 in Biology of Populus and its implications for management and conservation, edited by R. F. Stettler, H. D. Bradshaw, H. P. E and H. T. M. NRC Research Press, Ottawa, Canada.

Fang, L. C., H. L. Liu, S. Y. Wei, K. Keefover-Ring and T. M. Yin, 2018 High-density genetic map of *Populus deltoides* constructed by using specific length amplified fragment sequencing. Tree Genetics & Genomes 14: 10.

Fay, M. P., 2010 Two-sided exact tests and matching confidence intervals for discrete data. R Journal 2: 53–58.

Felsenstein, J., 1974 The evolutionary advantage of recombination. Genetics 78: 737–756.

Gaudet, M., V. Jorge, I. Paolucci, I. Beritognolo, G. S. Mugnozza et al., 2008 Genetic linkage maps of *Populus nigra* L. including AFLPs, SSRs, SNPs, and sex trait. Tree Genetics & Genomes 4: 25–36.

Gaut, B. S., S. I. Wright, C. Rizzon, J. Dvorak and L. K. Anderson, 2007 Recombination: an underappreciated factor in the evolution of plant genomes. Nature Reviews Genetics 8: 77–84.

Geraldes, A., C. A. Hefer, A. Capron, N. Kolosova, F. Martinez-Nunez et al., 2015 Recent Y chromosome divergence despite ancient origin of dioecy in poplars (*Populus*). Molecular Ecology 24: 3243–3256.

Gion, J. M., C. J. Hudson, I. Lesur, R. E. Vaillancourt, B. M. Potts et al., 2016 Genome-wide variation in recombination rate in *Eucalyptus*. Bmc Genomics 17: 12.

Giraut, L., M. Falque, J. Drouaud, L. Pereira, O. C. Martin et al., 2011 Genome-wide crossover distribution in *Arabidopsis thaliana* meiosis reveals sex-specific patterns along chromosomes. Plos Genetics 7: 15.

Grattapaglia, D., F. L. G. Bertolucci, R. Penchel and R. R. Sederoff, 1996 Genetic mapping of quantitative trait loci controlling growth and wood quality traits in *Eucalyptus grandis* using a maternal half-sib family and RAPD markers. Genetics 144: 1205–1214.

Grattapaglia, D., and M. D. V. Resende, 2011 Genomic selection in forest tree breeding. Tree Genetics & Genomes 7: 241–255.

Groover, A. T., C. G. Williams, M. E. Devey, J. M. Lee and D. B. Neale, 1995 Sex-related differences in meiotic recombination frequency in *Pinus taeda*. Journal of Heredity 86: 157–158.

Haenel, Q., T. G. Laurentino, M. Roesti and D. Berner, 2018 Meta-analysis of chromosome-scale crossover rate variation in eukaryotes and its significance to evolutionary genomics. Molecular Ecology 27: 2477–2497.

Harman-Ware, A. E., D. Macaya-Sanz, C. R. Abeyratne, C. Doeppke, K. Haiby et al., 2021 Accurate determination of genotypic variance of cell wall characteristics of a *Populus trichocarpa* pedigree using high-throughput pyrolysis-molecular beam mass spectrometry. Biotechnology for Biofuels 14: 15.

He, Y., M. H. Wang, S. Dukowic-Schulze, A. Zhou, C. L. Tiang et al., 2017 Genomic features shaping the landscape of meiotic double-strand-break hotspots in maize. Proceedings of the National Academy of Sciences of the United States of America 114: 12231–12236.

Heller, R., and H. Gur, 2011 False discovery rate controlling procedures for discrete tests. arXiv preprint arXiv:1112.4627.

Hey, J., 2004 What’s so hot about recombination hotspots? Plos Biology 2: 730–733.

Hiatt, E. N., and R. K. Dawe, 2003 Four loci on abnormal chromosome 10 contribute to meiotic drive in maize. Genetics 164: 699–709.

Hill, W. G., and A. Robertson, 1966 The effect of linkage on limits to artificial selection. Genetics Research 8: 269–294.

Hofmeister, B. T., J. Denkena, M. Colome-Tatche, Y. Shahryary, R. Hazarika et al., 2020 A genome assembly and the somatic genetic and epigenetic mutation rate in a wild long-lived perennial *Populus trichocarpa*. Genome Biology 21: 27.

Hunt, P. A., and T. J. Hassold, 2002 Sex matters in meiosis. Science 296: 2181–2183.

Isik, F., 2014 Genomic selection in forest tree breeding: the concept and an outlook to the future. New Forests 45: 379–401.

Jansen, R. C., J. L. Jannink and W. D. Beavis, 2003 Mapping quantitative trait loci in plant breeding populations: use of parental haplotype sharing. Crop Science 43: 829–834.

Kauppi, L., A. J. Jeffreys and S. Keeney, 2004 Where the crossovers are: recombination distributions in mammals. Nature Reviews Genetics 5: 413–424.

Kent, T. V., J. Uzunovic and S. I. Wright, 2017 Coevolution between transposable elements and recombination. Philosophical Transactions of the Royal Society B-Biological Sciences 372: 11.

Kianian, P. M. A., M. H. Wang, K. Simons, F. Ghavami, Y. He et al., 2018 High-resolution crossover mapping reveals similarities and differences of male and female recombination in maize. Nature Communications 9: 10.

Kim, G., A. P. L. Montalvao, B. Kersten, M. Fladung and N. A. Muller, 2021 The genetic basis of sex determination in *Populus* provides molecular markers across the genus and indicates convergent evolution. Silvae Genetica 70: 145–155.

Kong, A., G. Thorleifsson, D. F. Gudbjartsson, G. Masson, A. Sigurdsson et al., 2010 Fine-scale recombination rate differences between sexes, populations and individuals. Nature 467: 1099–1103.

Kuhn, M., 2008 Building predictive models in R using the caret package. Journal of Statistical Software 28: 1–26.

Lambing, C., F. C. H. Franklin and C. J. R. Wang, 2017 Understanding and Manipulating Meiotic Recombination in Plants. Plant Physiology 173: 1530–1542.

Lambing, C., A. J. Tock, S. D. Topp, K. Choi, P. C. Kuo et al., 2020 Interacting genomic landscapes of REC8-Cohesin, chromatin, and meiotic recombination in *Arabidopsis*. Plant Cell 32: 1218–1239.

Lange, J., S. Yamada, S. E. Tischfield, J. Pan, S. Kim et al., 2016 The landscape of mouse meiotic double-strand break formation, processing, and repair. Cell 167: 695–708.

Lenormand, T., 2003 The evolution of sex dimorphism in recombination. Genetics 163: 811–822.

Lenormand, T., and J. Dutheil, 2005 Recombination difference between sexes: A role for haploid selection. Plos Biology 3: 396–403.

Littrell, J., S. W. Tsaih, A. Baud, P. Rastas, L. Solberg-Woods et al., 2018 A high-resolution genetic map for the laboratory rat. G3-Genes Genomes Genetics 8: 2241–2248.

Lloyd, A., and E. Jenczewski, 2019 Modelling sex-specific crossover patterning in Arabidopsis. Genetics 211: 847–859.

Luo, C., X. Li, Q. H. Zhang and J. B. Yan, 2019 Single gametophyte sequencing reveals that crossover events differ between sexes in maize. Nature Communications 10: 8.

Marcais, G., and C. Kingsford, 2011 A fast, lock-free approach for efficient parallel counting of occurrences of k-mers. Bioinformatics 27: 764–770.

Margarido, G. R. A., A. P. Souza and A. A. F. Garcia, 2007 OneMap: software for genetic mapping in outcrossing species. Hereditas 144: 78–79.

Mezard, C., 2006 Meiotic recombination hotspots in plants. Biochemical Society Transactions 34: 531–534.

Moran, G. F., J. C. Bell and A. J. Hilliker, 1983 Greater meiotic recombination in male vs. female gametes in *Pinus radiata*. Journal of Heredity 74: 62–62.

Muller, H. J., 1964 The relation of recombination to mutational advance. Mutation Research 1: 2–9.

Muranty, H., V. Jorge, C. Bastien, C. Lepoittevin, L. Bouffier et al., 2014 Potential for marker-assisted selection for forest tree breeding: lessons from 20 years of MAS in crops. Tree Genetics & Genomes 10: 1491–1510.

Neale, D. B., and A. Kremer, 2011 Forest tree genomics: growing resources and applications. Nature Reviews Genetics 12: 111–122.

Pan, J., M. Sasaki, R. Kniewel, H. Murakami, H. G. Blitzblau et al., 2011 A hierarchical combination of factors shapes the genome-wide topography of yeast meiotic recombination initiation. Cell 144: 719–731.

Penalba, J. V., and J. B. W. Wolf, 2020 From molecules to populations: appreciating and estimating recombination rate variation. Nature Reviews Genetics 21: 476–492.

Petit, R. J., and A. Hampe, 2006 Some evolutionary consequences of being a tree, pp. 187–214 in Annual Review of Ecology Evolution and Systematics. Annual Reviews, Palo Alto.

Petkolv, P. M., K. W. Broman, J. P. Szatkiewicz and K. Paigen, 2007 Crossover interference underlies sex differences in recombination rates. Trends in Genetics 23: 539–542.

Phillips, D., G. Jenkins, M. Macaulay, C. Nibau, J. Wnetrzak et al., 2015 The effect of temperature on the male and female recombination landscape of barley. New Phytologist 208: 421–429.

Plomion, C., and D. M. Omalley, 1996 Recombination rate differences for pollen parents and seed parents in *Pinus pinaster*. Heredity 77: 341–350.

Porth, I., and Y. A. El-Kassaby, 2015 Using *Populus* as a lignocellulosic feedstock for bioethanol. Biotechnology Journal 10: 510–U214.

Pucholt, P., H. R. Hallingback and S. Berlin, 2017 Allelic incompatibility can explain female biased sex ratios in dioecious plants. Bmc Genomics 18: 12.

Quinlan, A. R., and I. M. Hall, 2010 BEDTools: a flexible suite of utilities for comparing genomic features. Bioinformatics 26: 841–842.

R Core Team, 2013 R: A language and environment for statistical computing, pp. R Foundation for statistical Computing, Vienna, Austria.

Raj, A., M. Stephens and J. K. Pritchard, 2014 fastSTRUCTURE: variational inference of population structure in large SNP data sets. Genetics 197: 573–U207.

Rodgers-Melnick, E., P. J. Bradbury, R. J. Elshire, J. C. Glaubitz, C. B. Acharya et al., 2015 Recombination in diverse maize is stable, predictable, and associated with genetic load. Proceedings of the National Academy of Sciences of the United States of America 112: 3823–3828.

Rodgers-Melnick, E., D. L. Vera, H. W. Bass and E. S. Buckler, 2016 Open chromatin reveals the functional maize genome. Proceedings of the National Academy of Sciences of the United States of America 113: E3177–E3184.

Rood, S. B., J. S. Campbell and T. Despins, 1986 Natural poplar hybrids from southern Alberta. I. Continuous variation for foliar characteristics. Canadian Journal of Botany 64: 1382–1388.

Rowan, B. A., D. Heavens, T. R. Feuerborn, A. J. Tock, I. R. Henderson et al., 2019 An ultra high-density *Arabidopsis thaliana* crossover map that refines the influences of structural variation and epigenetic features. Genetics 213: 771–787.

Saintenac, C., S. Faure, A. Remay, F. Choulet, C. Ravel et al., 2011 Variation in crossover rates across a 3-Mb contig of bread wheat (*Triticum aestivum*) reveals the presence of a meiotic recombination hotspot. Chromosoma 120: 185–198.

Sannigrahi, P., A. J. Ragauskas and G. A. Tuskan, 2010 Poplar as a feedstock for biofuels: A review of compositional characteristics. Biofuels Bioproducts & Biorefining-Biofpr 4: 209–226.

Sardell, J. M., and M. Kirkpatrick, 2020 Sex differences in the recombination landscape. American Naturalist 195: 361–379.

Segal, E., and J. Widom, 2009 Poly(dA:dT) tracts: major determinants of nucleosome organization. Current Opinion in Structural Biology 19: 65–71.

Shilo, S., C. Melamed-Bessudo, Y. Dorone, N. Barkai and A. A. Levy, 2015 DNA crossover motifs associated with epigenetic modifications delineate open chromatin regions in *Arabidopsis*. The Plant Cell 27: 2427–2436.

Sidhu, G. K., C. Fang, M. A. Olson, M. Falque, O. C. Martin et al., 2015 Recombination patterns in maize reveal limits to crossover homeostasis. Proceedings of the National Academy of Sciences of the United States of America 112: 15982–15987.

Silva - Junior, O. B., and D. Grattapaglia, 2015 Genome - wide patterns of recombination, linkage disequilibrium and nucleotide diversity from pooled resequencing and single nucleotide polymorphism genotyping unlock the evolutionary history of *Eucalyptus grandis*. New Phytologist 208: 830–845.

Slavov, G. T., S. P. DiFazio, J. Martin, W. Schackwitz, W. Muchero et al., 2012 Genome resequencing reveals multiscale geographic structure and extensive linkage disequilibrium in the forest tree *Populus trichocarpa*. New Phytologist 196: 713–725.

Slavov, G. T., S. Leonardi, J. Burczyk, W. T. Adams, S. H. Strauss et al., 2009 Extensive pollen flow in two ecologically contrasting populations of *Populus trichocarpa*. Molecular Ecology 18: 357–373.

Smagulova, F., I. V. Gregoretti, K. Brick, P. Khil, R. D. Camerini-Otero et al., 2011 Genome-wide analysis reveals novel molecular features of mouse recombination hotspots. Nature 472: 375–378.

Smit, A. F. A., R. Hubley and P. Green, 2013-2015 RepeatMasker Open-4.0, pp., http://www.repeatmasker.org.

Smukowski, C. S., and M. A. F. Noor, 2011 Recombination rate variation in closely related species. Heredity 107: 496–508.

Tiley, G. P., and G. Burleigh, 2015 The relationship of recombination rate, genome structure, and patterns of molecular evolution across angiosperms. Bmc Evolutionary Biology 15: 14.

Tuskan, G. A., S. DiFazio, S. Jansson, J. Bohlmann, I. Grigoriev et al., 2006 The genome of black cottonwood, *Populus trichocarpa* (Torr. & Gray). Science 313: 1596–1604.

Van Os, H., P. Stam, R. G. F. Visser and H. J. Van Eck, 2005 RECORD: a novel method for ordering loci on a genetic linkage map. Theoretical and Applied Genetics 112: 30–40.

Vining, K. J., K. R. Pomraning, L. J. Wilhelm, H. D. Priest, M. Pellegrini et al., 2012 Dynamic DNA cytosine methylation in the *Populus trichocarpa* genome: tissue-level variation and relationship to gene expression. Bmc Genomics 13: 19.

Wang, J. L., 2004 Sibship reconstruction from genetic data with typing errors. Genetics 166: 1963–1979.

Wang, M. C., L. Zhang, Z. Y. Zhang, M. M. Li, D. Y. Wang et al., 2020 Phylogenomics of the genus Populus reveals extensive interspecific gene flow and balancing selection. New Phytologist 225: 1370–1382.

Wang, Y. X., and G. P. Copenhaver, 2018 Meiotic recombination: mixing it up in plants, pp. 577–609 in Annual Review of Plant Biology, Vol 69, edited by S. S. Merchant. Annual Reviews, Palo Alto.

Wegrzyn, J. L., A. J. Eckert, M. Choi, J. M. Lee, B. J. Stanton et al., 2010 Association genetics of traits controlling lignin and cellulose biosynthesis in black cottonwood (*Populus trichocarpa*, Salicaceae) secondary xylem. New Phytologist 188: 515–532.

Wu, R., H. D. Bradshaw and R. F. Stettler, 1998 Developmental quantitative genetics of growth in *Populus*. Theoretical and Applied Genetics 97: 1110–1119.

Yelina, N. E., K. Choi, L. Chelysheva, M. Macaulay, B. de Snoo et al., 2012 Epigenetic remodeling of meiotic crossover frequency in *Arabidopsis thaliana* DNA methyltransferase mutants. Plos Genetics 8: 16.

Yin, T. M., S. P. DiFazio, L. E. Gunter, D. Riemenschneider and G. A. Tuskan, 2004 Large-scale heterospecific segregation distortion in *Populus* revealed by a dense genetic map. Theoretical and Applied Genetics 109: 451–463.

Yin, T. M., X. Y. Zhang, M. R. Huang, M. X. Wang, Q. Zhuge et al., 2002 Molecular linkage maps of the *Populus* genome. Genome 45: 541–555.

Zelkowski, M., M. A. Olson, M. H. Wang and W. Pawlowski, 2019 Diversity and determinants of meiotic recombination landscapes. Trends in Genetics 35: 359–370.

Zhou, R., D. Macaya-Sanz, J. Schmutz, J. W. Jenkins, G. A. Tuskan et al., 2020 Sequencing and analysis of the sex determination region of *Populus trichocarpa*. Genes 11: 20.

