## Supplementary Methods for "High resolution mapping reveals hotspots and sex-biased recombination in *Populus trichocarpa*"

**Discrete wavelet analysis for identifying the scales at which sex-based CO count differences are most prominent:**

Patterns of sex-based differences observed at the chromosome scale could manifest due to underlying finer scale variation in CO rate between sexes. Here, the appropriate window size of comparison in CO counts was identified using a ‘wavelet’ analysis. A wavelet transformation decomposes the longitudinal CO count observations for each chromosome into a series of coefficients that explains the contribution at a range of different scales. Wavelet analysis is a signal processing technique that has been used in a number of other studies that analyzed genomic datasets, to unravel fine-scale spatial configuration of recombination across chromosomes (Spencer *et al.* 2006; Chan *et al.* 2012; Bherer *et al.* 2017; Weighill *et al.* 2018). The main advantage of this method is that it enables the detection of optimum window size as a result of the analysis rather than having to pre-select a window size. The input data for our analysis consisted of a numeric vector of cumulative CO counts within 30 kb bins across each chromosome for female and male groups separately (chromosome-wide signal for each sex). This comparison is justified given COs are contributed by an equal number of gametes by each sex. Decomposition of input signals was carried out using a maximum overlap discrete wavelet transformation (MODWT) with a ‘Haar’ mother wavelet. This method decomposed the variance in the original CO count signal into multiple orthogonal dyadic scales for each sex and each chromosome, and was implemented using the WMTSA R-package (Percival and Walden 2000). The lowest scale used here was 30kb and the highest was 3.84 Mb (representing a dyadic scale of 2^0^ to 2^7^, hereafter referred to as d0 to d7). Scales higher than these were not considered given the potential redundancy with whole chromosome scale analysis.

The wavelet analysis provided a decomposition of the total variance at each of the contributing scales for each sex and each chromosome (“wavVar” function in WMTSA). As expected, the lower most scales contributed much of the variance. Given the resolution of CO regions in our dataset, at this scale CO count data tend to be noisier, explaining the relatively high variance. In order to determine which dyadic scales were most sensitive to sex differences in each chromosome, a sex averaged (null) distribution of variance at each scale was estimated. Briefly, preserving the spatial configuration of the CO counts signal, common parent of each half-sib family was assigned to either male or female groupings randomly, resulting in equal number of parents in each group. The cumulative CO counts within 30 kb windows were calculated for each of the nominal groups and for each permutation (1000 permutations carried out), to obtain a sex indifferent null dataset. The empirical null distribution was used to obtain a p-value for the observed variances at a given scale for male and female groups (Figure S5). The 960 kb (d5) was selected as the best scale, as starting at this scale, majority of chromosomes in our dataset showed more extreme CO count variance (p ≥ 0.9 or p ≤ 0.1) for either observed male or female groups. In general, for any given cutoff we used (p=0.2 to 0.05) best window size scaled from d4 to d6 (corresponding to 480 kb to 1.9 Mb).

In order to visualize patterns of similarity in CO counts between sexes across multiple different scales across a chromosome, we implemented a continuous wavelet transformation (CWT) for each dataset using a ‘derivative of Gaussian wavelet’ basis (**Figure 2**). This transformation produces a set of wavelet coefficients for each genomic location that are normalized to reflect a variance of 1 at each scale. The CWT coefficient landscape can be plotted as a power spectrum for each scale that reflects CO count signal intensity at each genomic location at each scale. This power spectrum plot summarizes the CO count landscape and helps visualize overarching patterns between the two sexes. For broader scales, chromosome specific CO count patterns emerge where they are most noticeable between the scales d4 to d6 as expected based on our previous analysis, while in most cases differences between sexes at the finest scales remained indiscernible (Figure S6). However due to the highly redundant nature of scales used in CWT, it is presented here for mere illustration of broad-scale patterns between sexes and none of the statistical evaluations were based on them.

**REFERENCES:**

Bherer, C., C. L. Campbell and A. Auton, 2017 Refined genetic maps reveal sexual dimorphism in human meiotic recombination at multiple scales. Nature Communications 8**:** 9.

Chan, A. H., P. A. Jenkins and Y. S. Song, 2012 Genome-wide fine-scale recombination rate variation in *Drosophila melanogaster*. Plos Genetics 8**:** 28.

Percival, D. B., and A. T. Walden, 2000 *Wavelet methods for time series analysis (WMTSA)*. Cambridge University Press.

Spencer, C. C. A., P. Deloukas, S. Hunt, J. Mullikin, S. Myers *et al.*, 2006 The influence of recombination on human genetic diversity. Plos Genetics 2**:** 1375-1385.

Weighill, D., P. Jones, M. Shah, P. Ranjan, W. Muchero *et al.*, 2018 Pleiotropic and epistatic network-based discovery: integrated networks for target gene discovery. Frontiers in Energy Research 6**:** 20.
