## Supplementary Figures for "High resolution mapping reveals hotspots and sex-biased recombination in *Populus trichocarpa*"

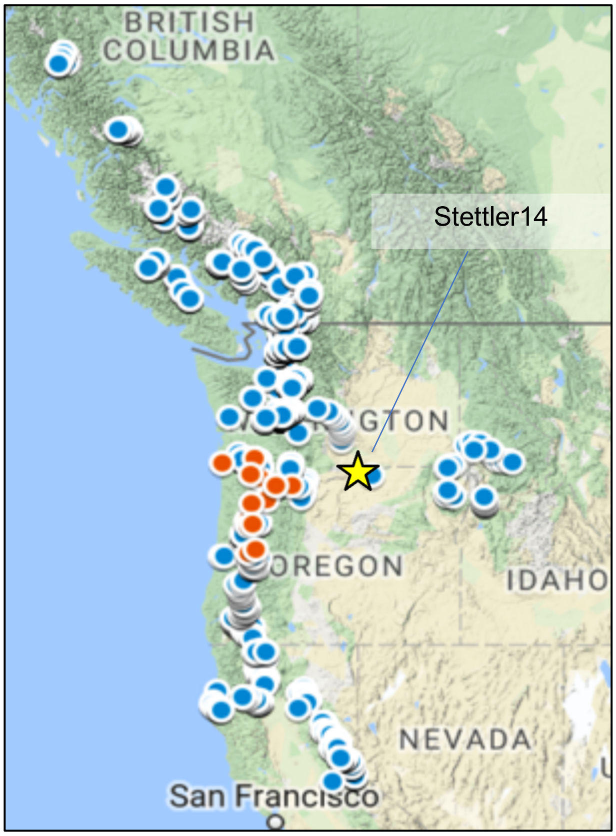


**Figure S1: Geographical location of parental clones in the 7x7 cross.**: The mapping population we use here is a subset of the GWAS population (WEGRZYN *et al.* 2010) in western USA and Canada which include individuals from the whole natural range Washington, Oregon and California in The USA and British Columbia, Canada. Circles indicate where the GWAS clones were collected and the 14 red circles indicate parents of the 7x7 trial. The yellow star marks the location of the *Populus trichocarpa* male collected from the Mount Hood region used as the reference genome (Stettler-14) in this study.

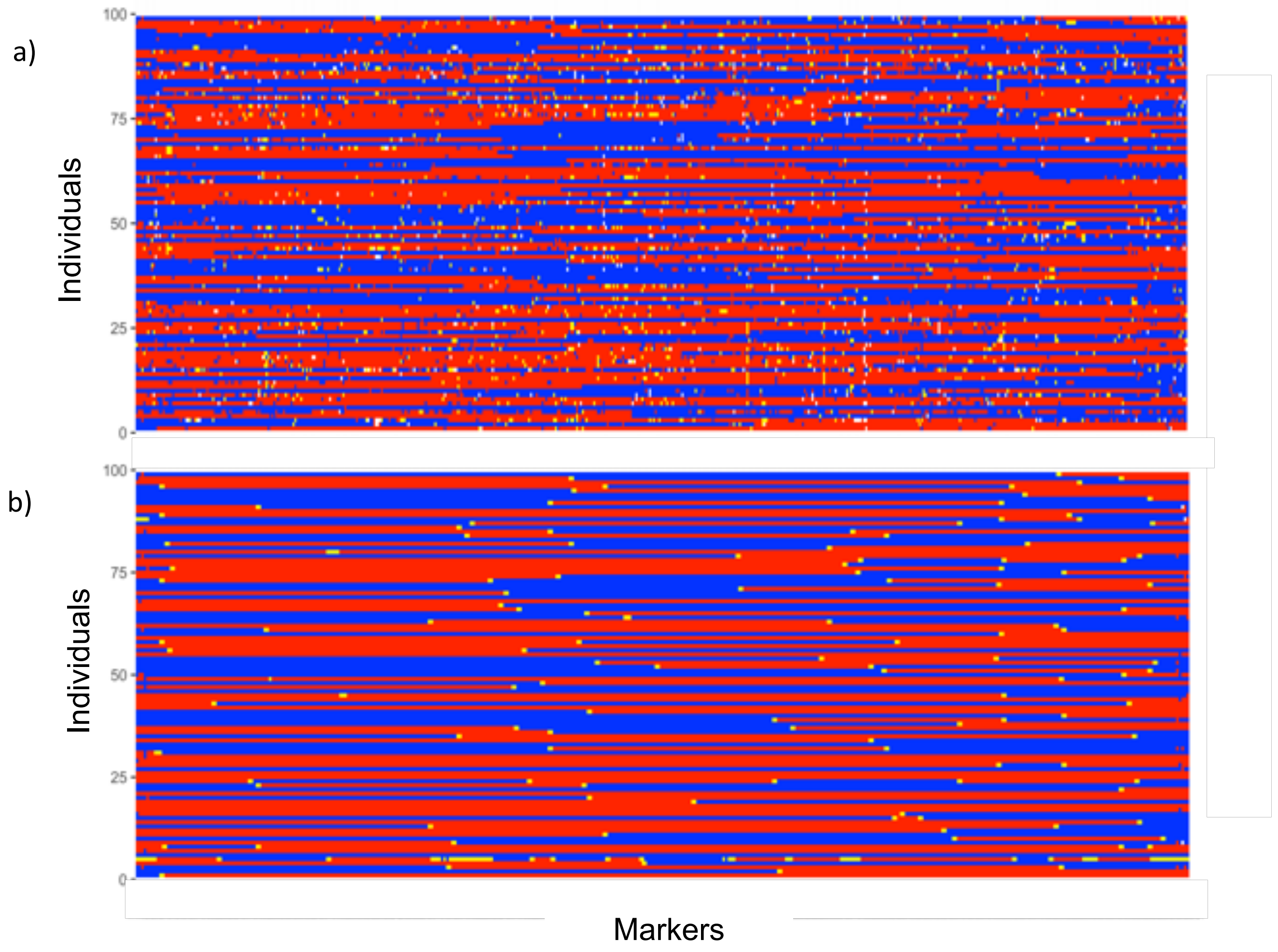


**Figure S2: Haplotype phasing and imputation. a)** Observed and **b)** imputed haplotypes, for offspring of the half-sib family of female parent GW-1863 on LG-III. Offspring were phased using trio genotype information. red or blue lines represent each of the maternal haplotypes. Missing genotypes and Mendelian violations are marked in yellow. This process was completed for all 19 LGs for each of the 14 parents.

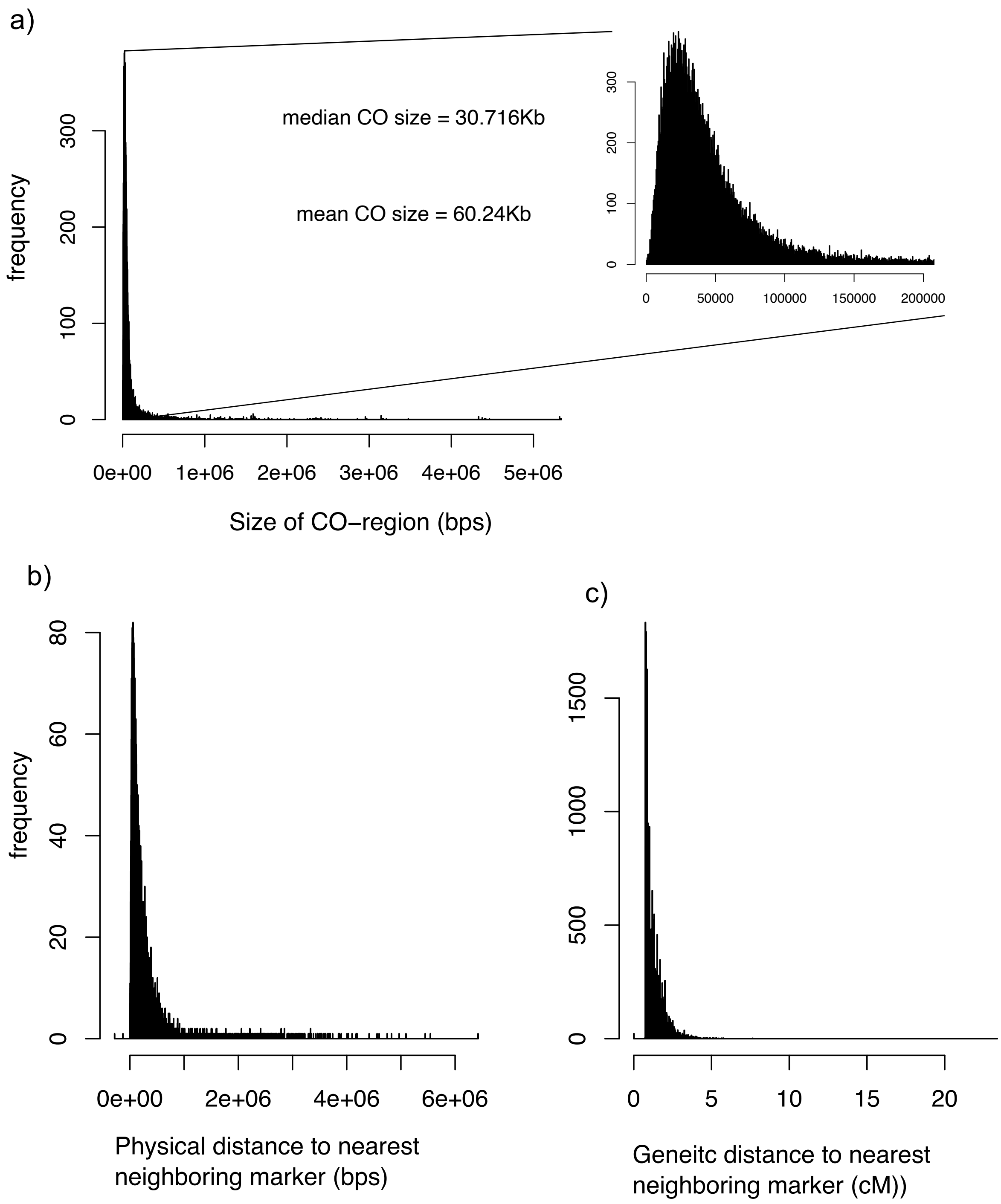


**Figure S3:** **(a)** CO region size distribution for all 19 LGs for all half-sib families; **(b)** median physical distance between neighboring markers in all genetic maps; **(c)** median genetic map distance between neighboring markers for all genetic maps.

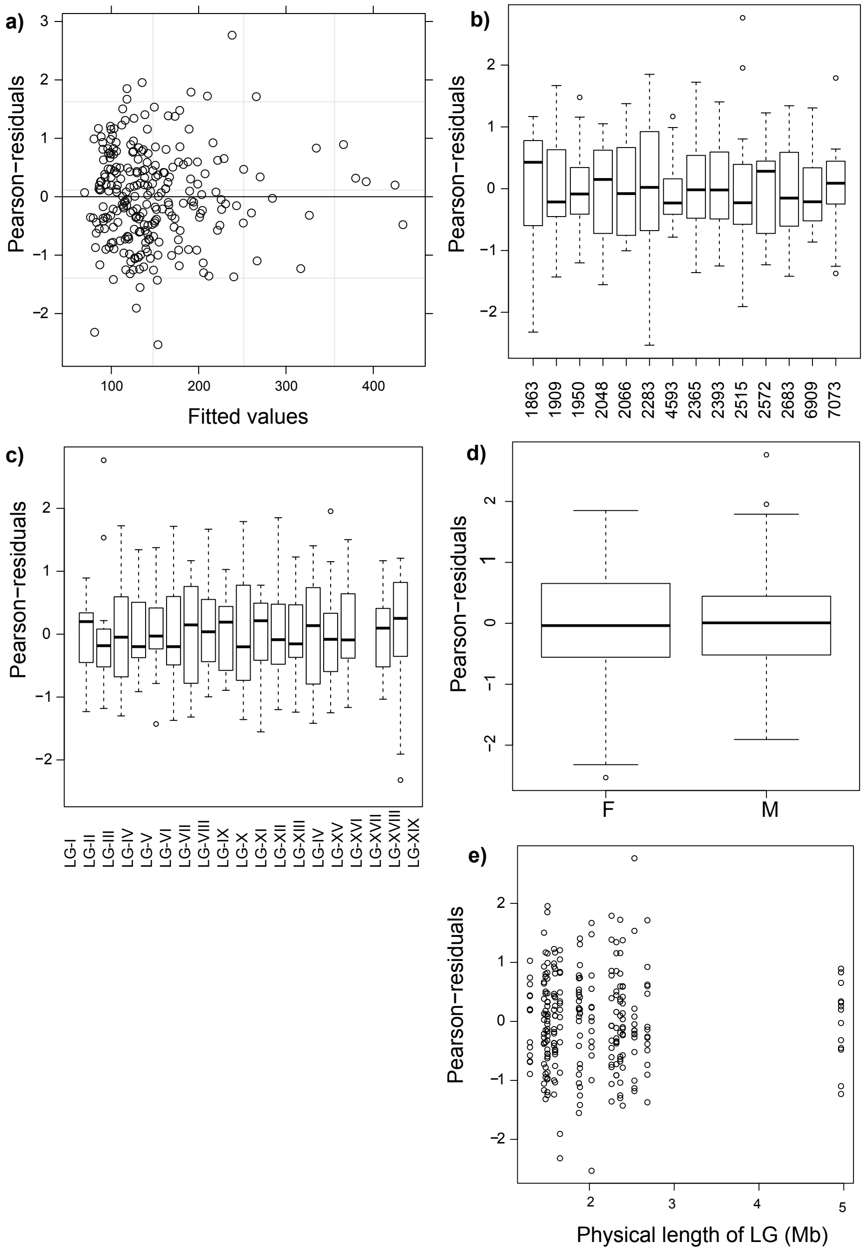


**Figure S4: Statistical model fit for GLMM analyzing chromosome scale sex-based differences in CO counts:** The distribution of Pearson residuals derived after fitting the GLMM with a log-link function did not show any significant trends across fitted values **(a)** or any of the explanatory variables **(b & e)** such as half-sib family identity, LG identity, sex or physical length of the chromosome

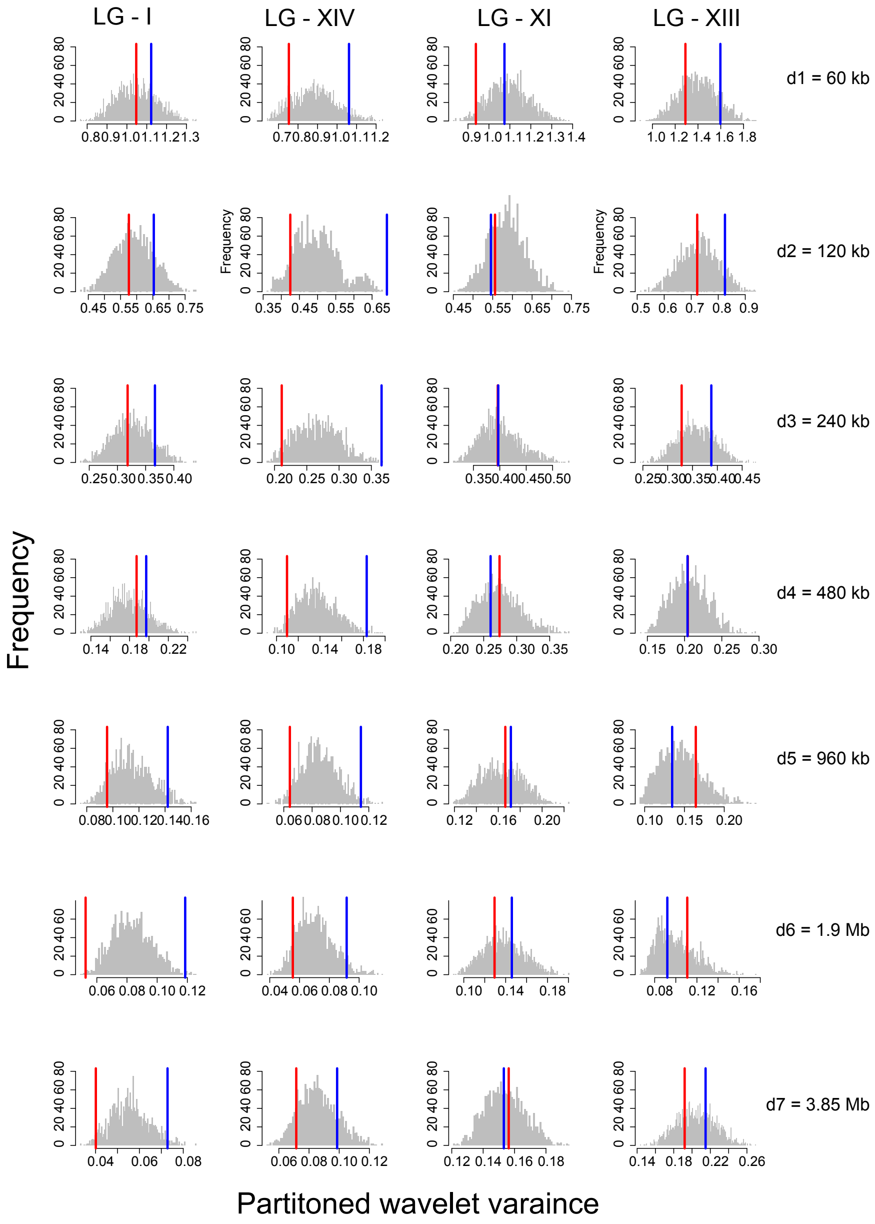


**Figure S5: Partitioned wavelet variance.** Histograms (grey shading) represent null distribution of partitioned wavelet variance at each scale from 60 kb through 3.85 Mb (in rows), for four selected chromosomes (in columns). Null distributions were derived by randomly assigning parents to nominal groups of sex while preserving spatial structure of windows within each chromosome. The red and blue vertical lines represent the observed wavelet variance at each scale for female and male groups respectively. Significant separation between red and blue lines from 60 kb through 3.85 Mb are characteristic of LGs in which heterochiasmy is present such as LG-I, LG-XIV, while others such as LG-XI and LG-XIII show no such trend and do not show differences between sexes.

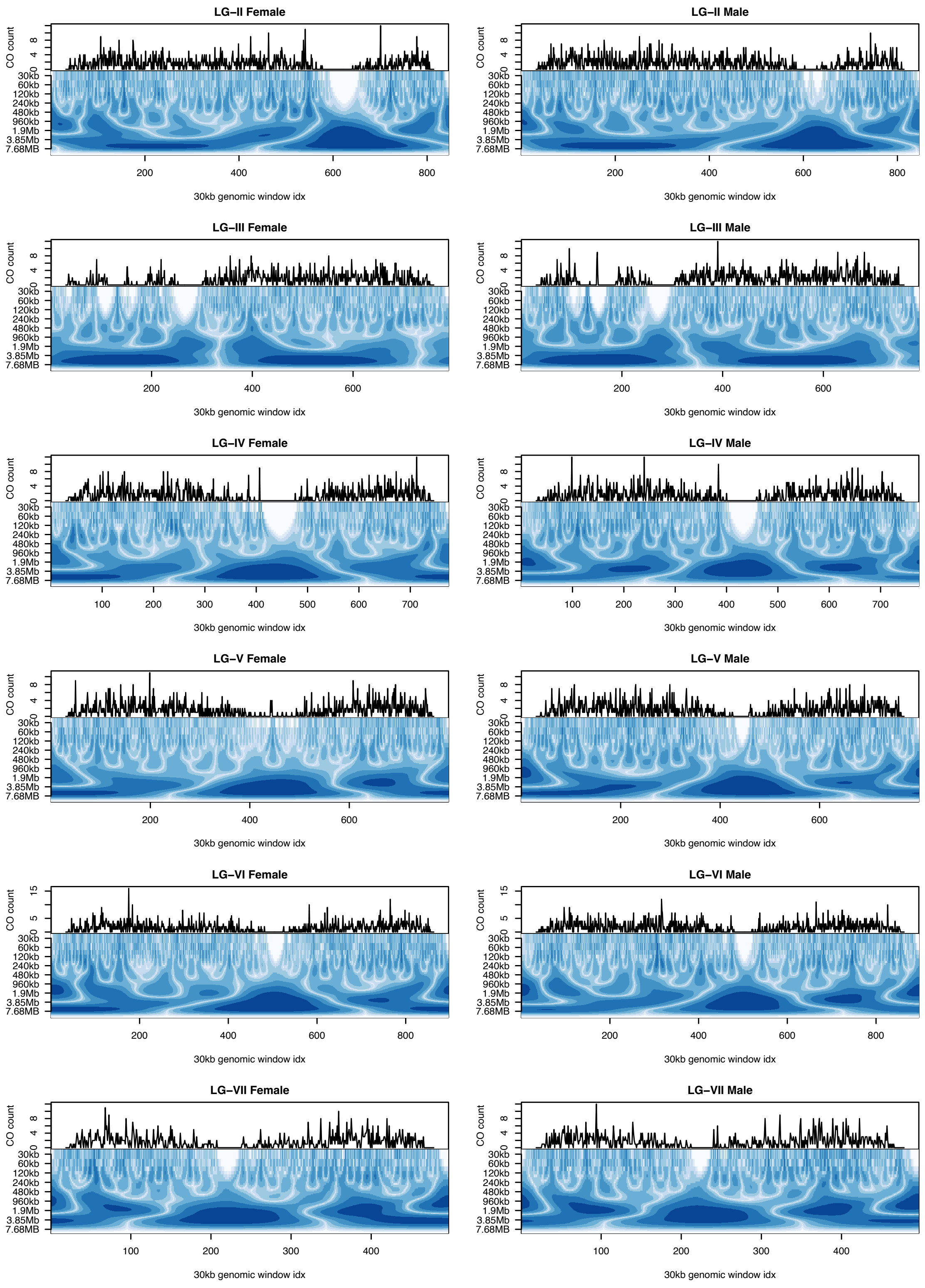
LG–II Female
LG–II Male
LG–III Female
LG–III Male
LG–IV Female
LG–IV Male
LG–V Female
LG–V Male
LG–VI Female
LG–VI Male
LG–VII Female
LG–VII Male
CO count
30kb
60kb
120kb
240kb
480kb
960kb
1.9Mb
3.85Mb
7.68MB
30kb genomic window idx

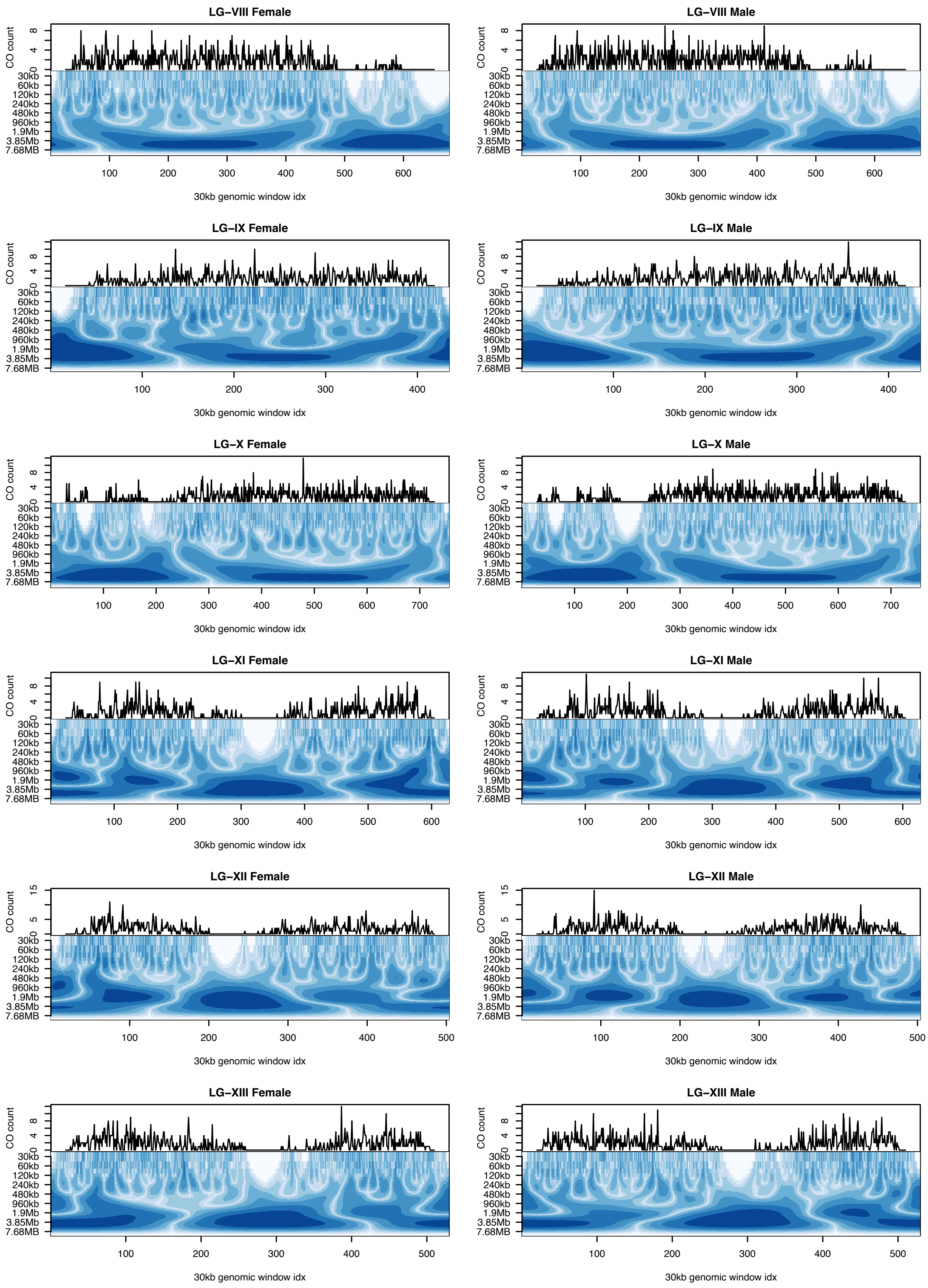

LG–VIII Female
LG–VIII Male
LG–IX Female
LG–IX Male
LG–X Female
LG–X Male
LG–XI Female
LG–XI Male
LG–XII Female
LG–XII Male
LG–XIII Female
LG–XIII Male
CO count
30kb
60kb
120kb
240kb
480kb
960kb
1.9Mb
3.85Mb
7.68MB
30kb genomic window idx

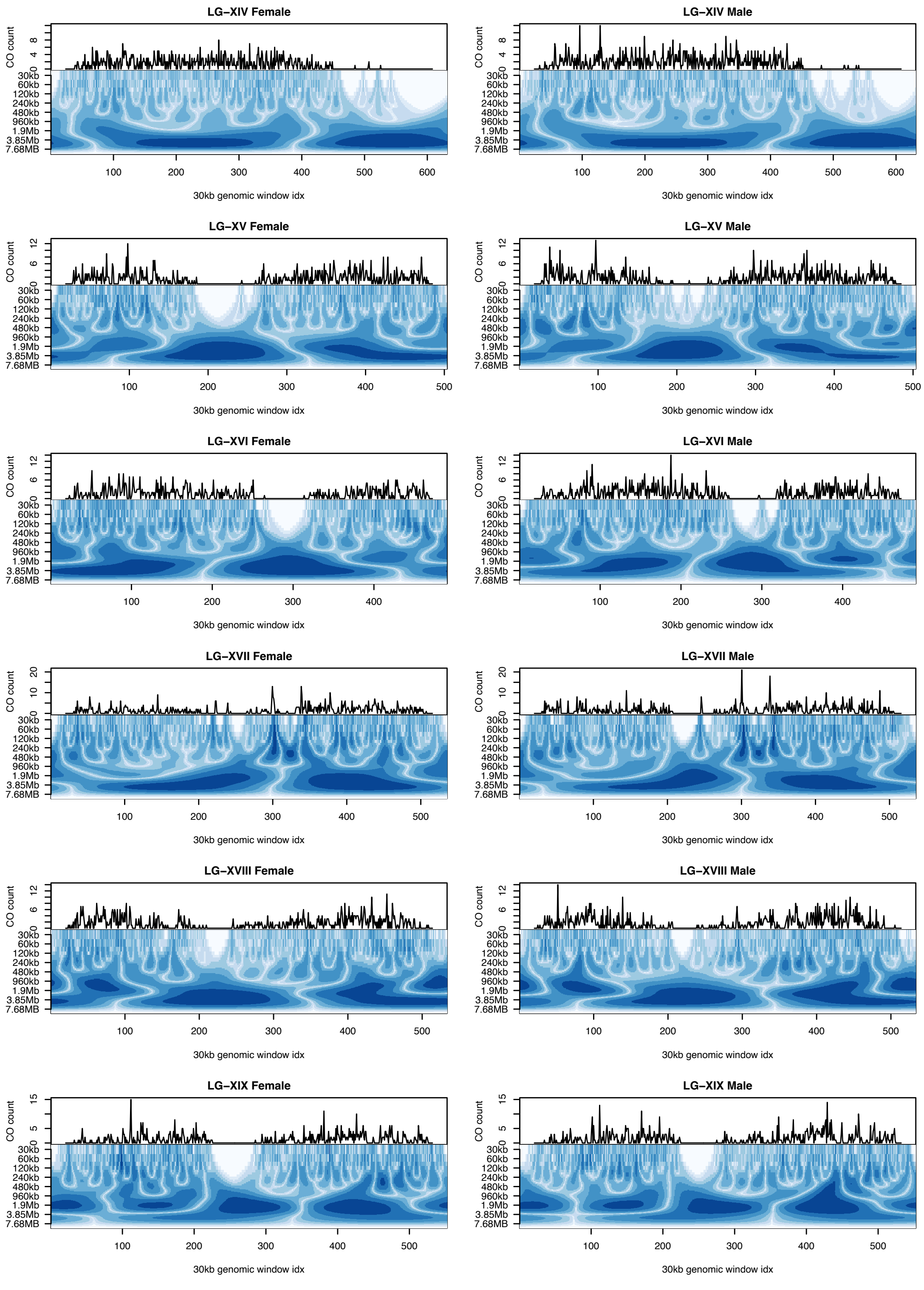


**Figure S6: Power spectrum of continuous wavelet transformation (CWT).** LG-II through LG-XIX (rows) are shown for male and female (columns) cumulative CO counts at each scale ranging from 30 kb to 7.68 MB.

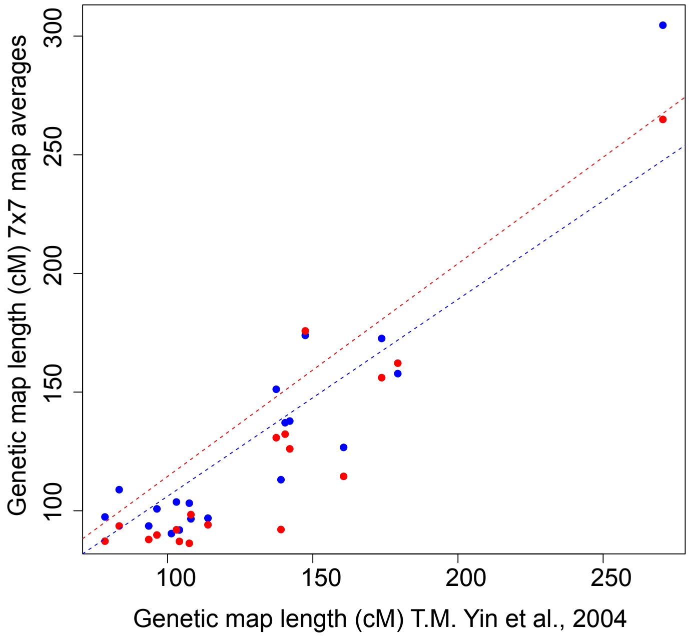


**Figure S7:** A high correlation was observed between genetic map averages produced in this study in comparison to similar maps from an independent study using SSR markers (YIN *et al.* 2004) Both male averaged (blue) and female averaged (red) map sizes showed high correlation (median female maps: $r = 0.92$, $p= 1.709 \times 10^{-8}$; median male maps: 0.93, $p= 5.868 \times 10^{-9}$). Blue and red dashed lines represent the linear regression fit for male and female median genetic maps for 19 linkage groups respectively.

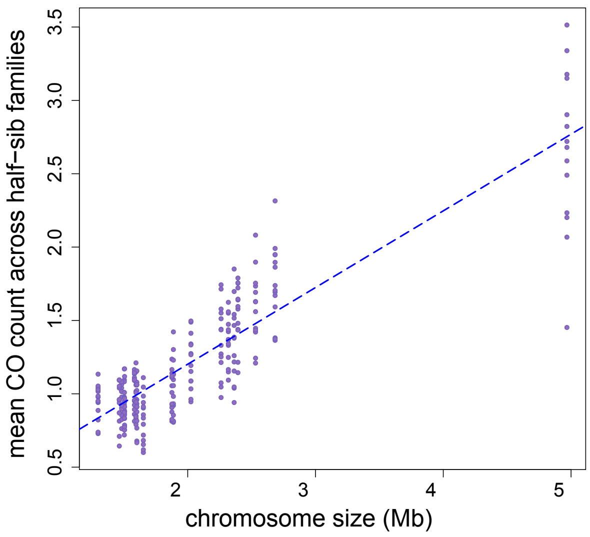


**Figure S8: Regression of mean CO count vs. physical length for a given chromosome.** Dots represent the mean CO count for a given half-sib family for a given chromosome. The dashed blue line represents the linear regression fit. Physical length of the chromosome explained majority of the variation in mean CO counts ($R^2$ = 0.77, p=2.2 x$10^{-16}$). There is considerable variance between half-sib families for a given chromosome and the magnitude increases as the size of the chromosome increases.

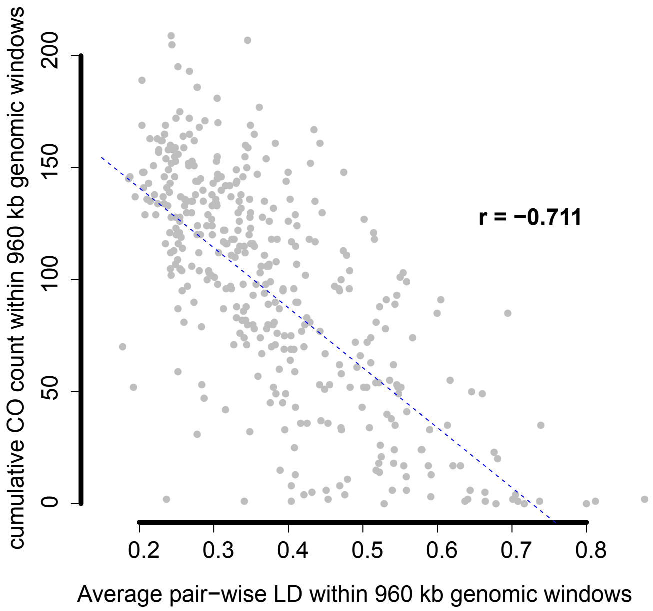


**Figure S9: Comparison of pairwise LD and cumulative CO counts:** A comparison of pairwise LD and cumulative CO counts across genomic windows of size 960 kb. A strong negative correlation was observed between cumulative CO counts and mean pairwise LD ($r_{Spearman}$= -0.711, p=2.2 x $10^{-16}$).

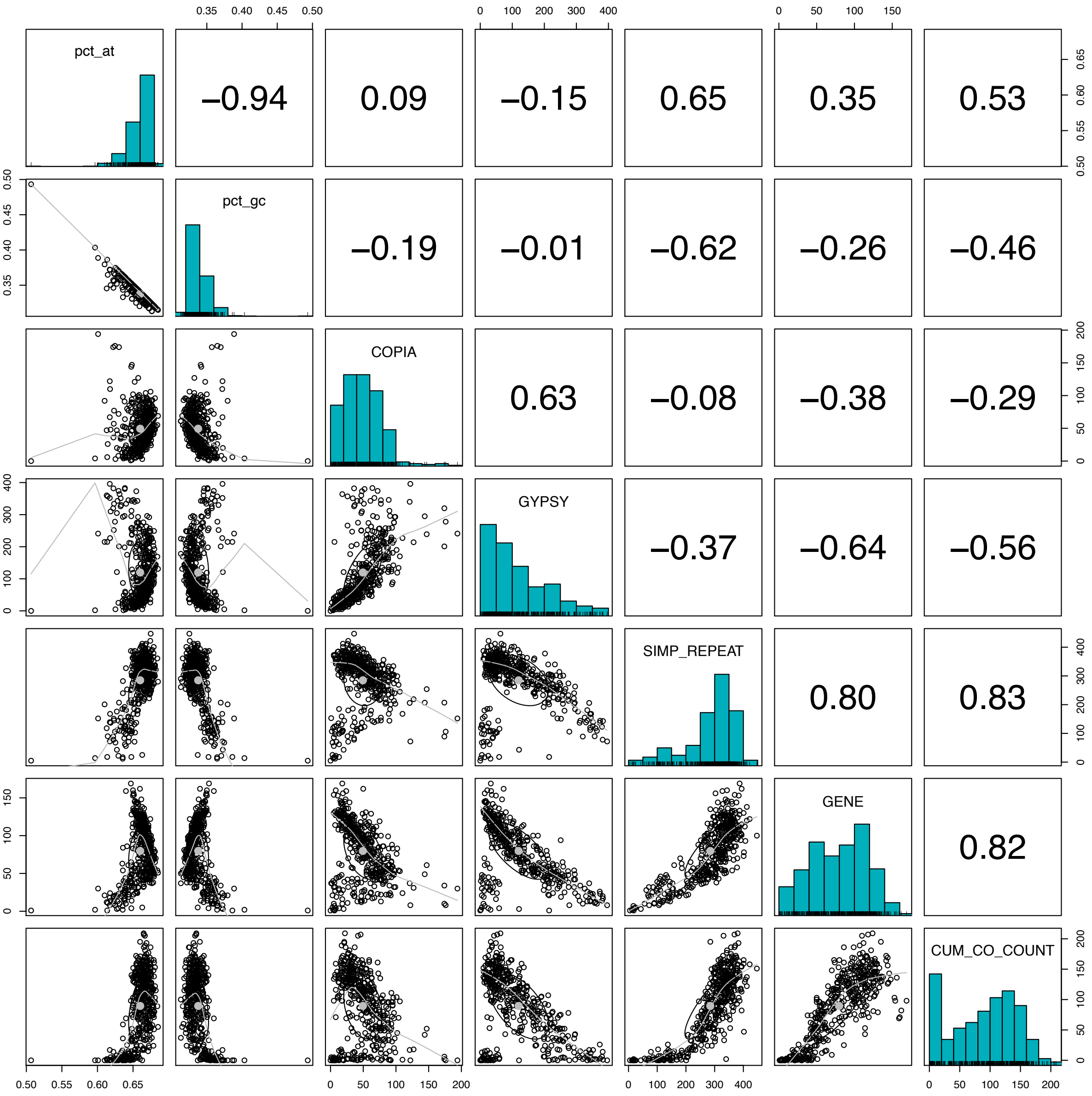


**Figure S10: CO pattern prediction using genomic correlates at 1Mb scale**. This shows the pair-wise relationship between various explanatory genomic features used to predict CO count within a given genomic region of size 1Mbps. There is high correlation among LTR families, as well as among LTRs, gene content, and GC content. High multi-collinearity is expected when these variables are used in linear models.

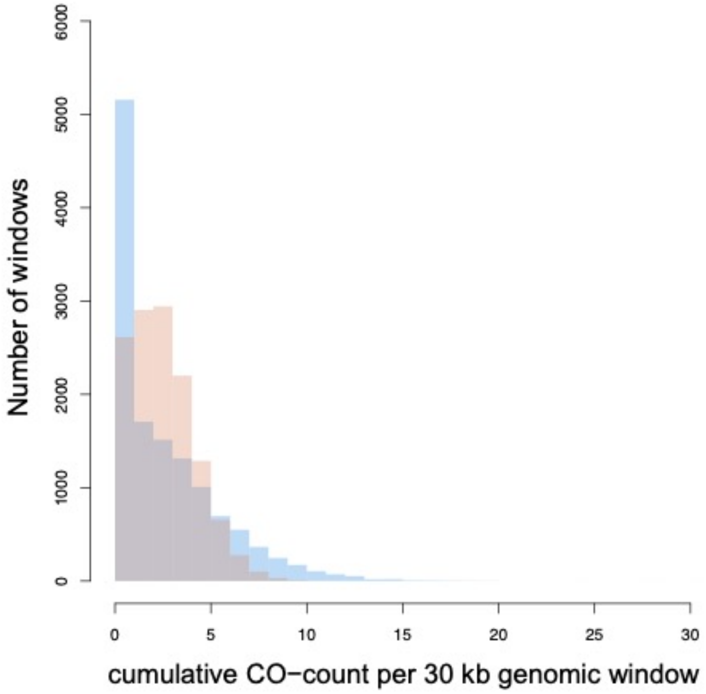


**Figure S11:** Empirical (blue) vs. null (orange) distributions of CO counts per 30 kb non-overlapping window size across the genome. The null distribution assumes that COs are distributed independently and identically across the genome, conforming to a theoretical Poisson distribution ($\lambda_{cumulative}$= 2.98). Observed CO counts per window are enriched for zero counts most likely due to windows that overlap centromeric span while longer than expected tail region indicate CO hotspot candidates

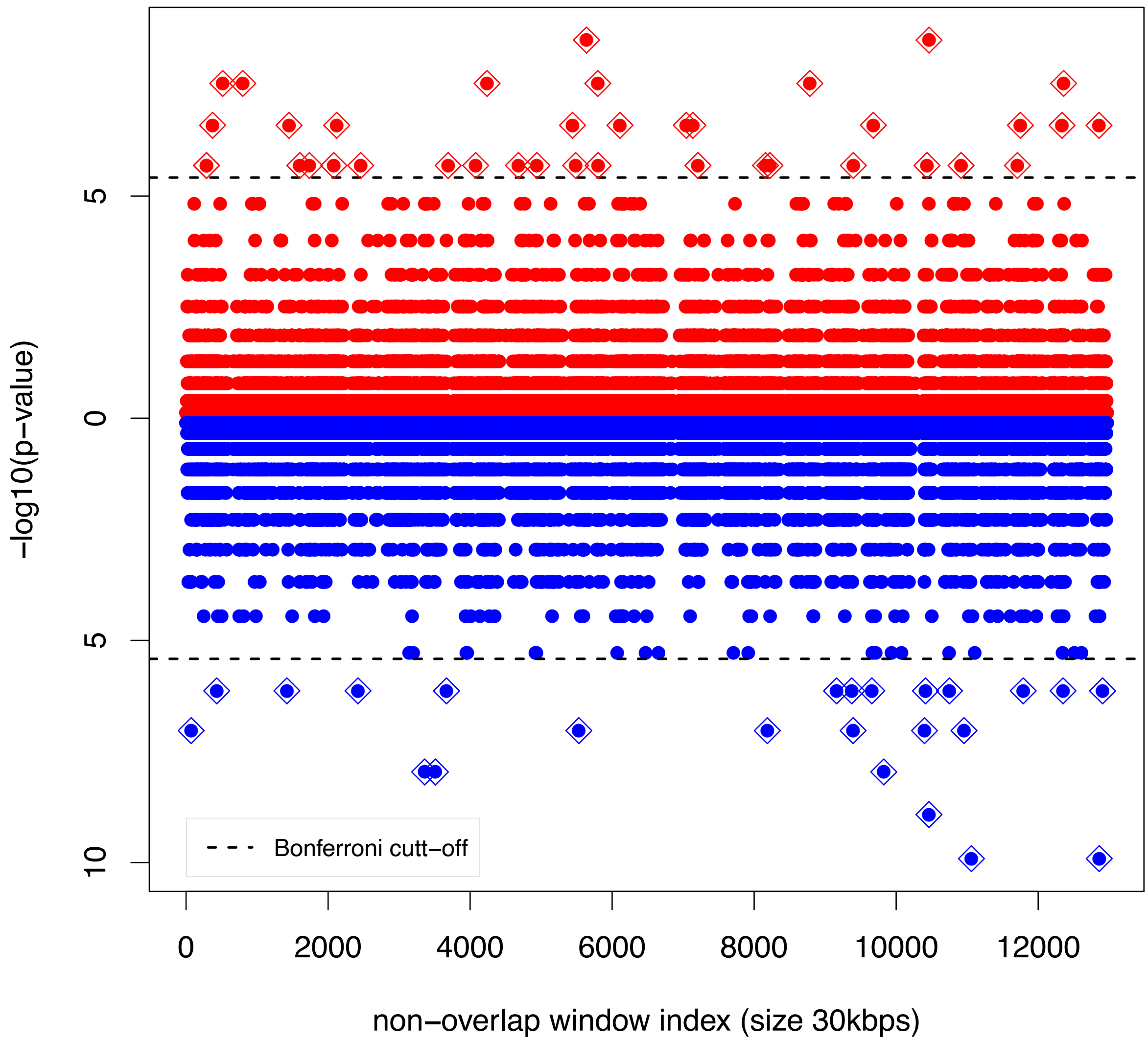


**Figure S12: CO hotspots for males and females.** Genome-wide hotspots were evaluated separately for males (blue) and females (red) with adjusted CO-counts per window for males and females and 38 and 24 CO hotspot windows were identified respectively ($\lambda_{female}$ =1.38; $\lambda_{male}$ =1.54; $FWER_{Bonferroni} \leq 0.05$). 21 and 11 windows overlapped with genomic scale cumulative CO-count hotspots for females and males respectively. Female and male hotspots were within the general vicinity of each other.

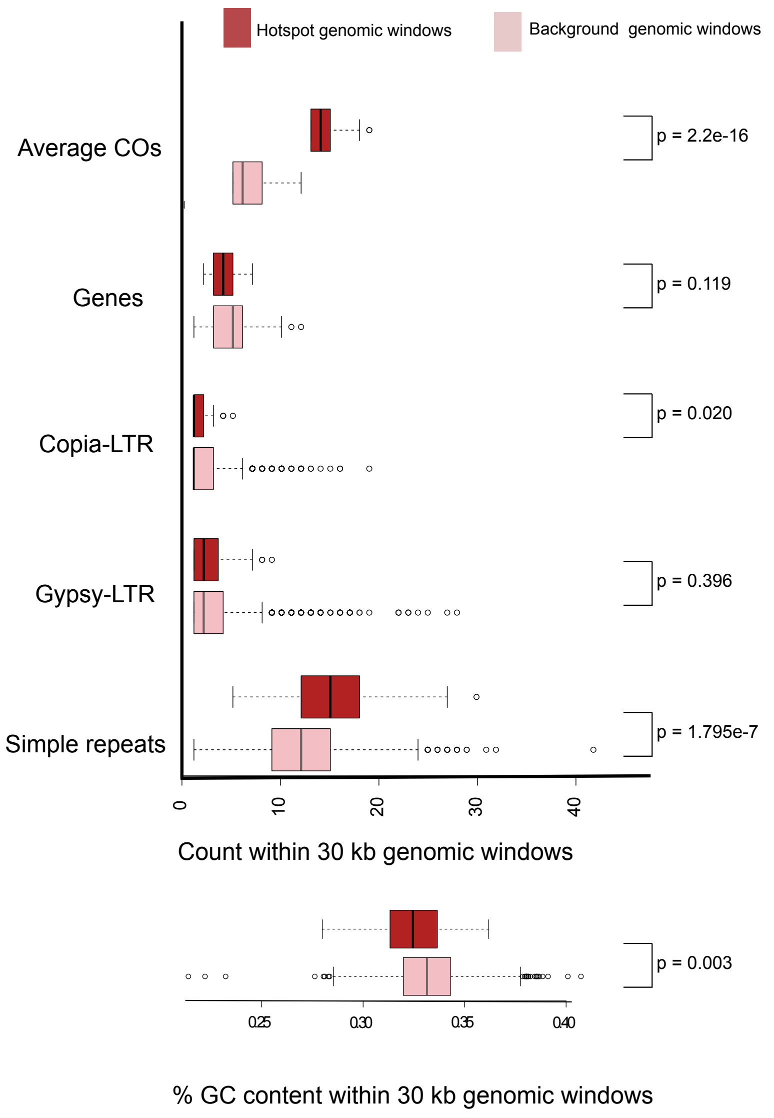


**Figure S13: Genomic correlates that influence localized CO patterns.** Genomic correlates associated with CO hotspots, in 30 kb genomic windows that showed significantly elevated CO counts (hotspots) with windows with background levels of CO counts.
